# The linear ubiquitin chain assembly complex LUBAC generates heterotypic ubiquitin chains

**DOI:** 10.1101/2020.05.27.117952

**Authors:** Alan Rodriguez Carvajal, Carlos Gomez Diaz, Antonia Vogel, Adar Sonn-Segev, Katrin Schodl, Luiza Deszcz, Zsuzsanna Orban-Nemeth, Shinji Sakamoto, Karl Mechtler, Philipp Kukura, Tim Clausen, David Haselbach, Fumiyo Ikeda

**Affiliations:** Institute of Molecular Biotechnology of the Austrian Academy of Sciences (IMBA), Vienna BioCenter (VBC), Dr. Bohr-Gasse 3, 1030 Vienna, Austria; Research Institute of Molecular Pathology (IMP), Vienna BioCenter (VBC), Campus-Vienna-Biocenter 1, 1030 Vienna, Austria; Department of Chemistry, University of Oxford, Chemistry Research Laboratory, 12 Mansfield Road, Oxford, OX1 3TA, UK; Pharmaceutical Frontier Research Labs, JT Inc., 13-2, Fukuura 1-chome, Kanazawa-ku, Yokohama, 236-0004, Japan; Medical Institute of Bioregulation (MIB), Kyushu University, 3-1-1 Maidashi, Higashiku, Fukuoka 812-8582, Japan

**Keywords:** E3 ubiquitin ligase, Ester-bond linkage, HOIL-1L, RBR, Ubiquitin

## Abstract

The linear ubiquitin chain assembly complex (LUBAC) is the only known ubiquitin ligase that generates linear/Met1-linked ubiquitin chains. One of the LUBAC components, HOIL-1L, was recently shown to catalyse oxyester bond formation between the C-terminus of ubiquitin and some substrates. However, oxyester bond formation in the context of LUBAC has not been directly observed. We present the first 3D reconstruction of LUBAC obtained by electron microscopy and report its generation of heterotypic ubiquitin chains containing linear linkages with oxyester-linked branches. We found that addition of the oxyester-bound branches depends on HOIL-1L catalytic activity. We suggest a coordinated ubiquitin relay mechanism between the HOIP and HOIL-1L ligases supported by cross-linking mass spectrometry data, which show proximity between the catalytic RBR domains. Mutations in the linear ubiquitin chain-binding NZF domain of HOIL-1L reduces chain branching confirming its role in the process. In cells, these heterotypic chains were induced by TNF. In conclusion, we demonstrate that LUBAC assembles heterotypic ubiquitin chains with linear and oxyester-linked branches by the concerted action of HOIP and HOIL-1L.

## Introduction

Posttranslational modification of substrates with ubiquitin (ubiquitination) regulates a wide variety of biological functions. Ubiquitin forms chains via its seven internal Lys residues (Lys6, Lys11, Lys27, Lys29, Lys33, Lys48, and Lys63) through an isopeptide bond, or via Met1 through a peptide bond (Komander & Rape, 2012). The different ubiquitin chain types contribute to determine the fate of the substrate and biological outcomes regulated.

For substrate modification, ubiquitin can be conjugated through isopeptide bonds to Lys residues, thioester bonds formed with the side chain of Cys residues, or oxyester bonds formed with side chains of Ser and Thr residues (Carvalho et al., 2007, McClellan et al., 2019, McDowell & Philpott, 2013, Vosper et al., 2009, Wang et al., 2007, Williams et al., 2007). In addition, some bacteria have evolved a ubiquitination mechanism, carried out by proteins of the SidE effector family, that results in phosphoribosyl-linked ubiquitin conjugated to Ser residues of the protein substrate (Bhogaraju et al., 2016, Qiu et al., 2016, Shin et al., 2020). Ubiquitin also contains seven Thr (Thr7, Thr9, Thr12, Thr14, Thr22, Thr55, and Thr66) and three Ser (Ser20, Ser57, and Ser65) residues that could potentially act as sites for chain formation. Recently, ubiquitin polymerization through Ser or Thr residues by a mammalian RBR-type ubiquitin ligase, Heme-Oxidized IRP2 Ubiquitin Ligase 1 (HOIL-1L) has been described (Kelsall et al., 2019). The E3 ligase MYCBP2 was also shown to conjugate the ubiquitin to Thr residues through an ester bond (Pao et al., 2018). Otherwise, oxyester bonds have only been observed between ubiquitin and other substrate proteins (McClellan et al., 2019, McDowell & Philpott, 2013).

HOIL-1L is a component of the Linear Ubiquitin chain Assembly Complex (LUBAC). LUBAC is thus far the only known E3 ubiquitin ligase complex that assembles linear/Met1-linked ubiquitin chains. LUBAC consists of two RING-in between-RING (RBR)-containing proteins: HOIL-1-Interacting Protein (HOIP) and HOIL-1L (Kirisako et al., 2006). LUBAC additionally contains the accessory protein Shank-Associated RH Domain-Interacting Protein (SHARPIN) (Gerlach et al., 2011, Ikeda et al., 2011, Tokunaga et al., 2011). HOIP has a catalytic center in its RING2 domain responsible for assembly of linear ubiquitin chains, while HOIL-1L and SHARPIN have been recognized as accessory proteins for the process. It is more recent that HOIL-1L has been shown to catalyse ubiquitination (Pao et al., 2018, Smit et al., 2013, Stieglitz et al., 2012).Linear ubiquitin chains and the three LUBAC components are essential components in biological functions including immune signalling (Gerlach et al., 2011, Ikeda, 2015, Iwai & Tokunaga, 2009, Rahighi et al., 2009, Rittinger & Ikeda, 2017, Tokunaga et al., 2009), development in mice (Fujita et al., 2018, Peltzer et al., 2018, Peltzer et al., 2014), protein quality control (van Well et al., 2019), Wnt signalling (Rivkin et al., 2013), and xenophagy (Noad et al., 2017, van Wijk et al., 2017). Therefore, it is important to understand how LUBAC assembles ubiquitin chains at the molecular level including how the catalytic activity of HOIL-1L contributes to the process.

In a recent study, Kelsall et al demonstrated that recombinant HOIL-1L can polymerise ubiquitin via oxyester bonds on Ser and Thr. They also showed that a HOIL-1L C458S mutation in which a predicted ubiquitin-loading site is mutated results in reduction of oxyester-linked ubiquitination signals in cells suggesting their dependency on HOIL-1L (Kelsall et al., 2019). Moreover, Fuseya et al. recently demonstrated that HOIL-1L catalytic activity negatively regulates the TNF signalling cascade (Fuseya et al., 2020). However, understanding the precise mechanisms regulating this atypical form of ubiquitination and whether HOIL-1L as a part of LUBAC mediates this biochemical activity remains largely unresolved.

## Results

### Reconstitution and 3D reconstruction of the LUBAC complex

We first set out to purify high quality recombinant LUBAC for structural characterization and biochemical investigation. Purifications of the three LUBAC components expressed individually in *E. coli* consistently gave low yields and isolated proteins were co-purified with several contaminants; this was particularly severe in purifications of full length HOIP (Figure 1A). Given that HOIP is destabilized in cells lacking SHARPIN or HOIL-1L (Fujita et al., 2018, Gerlach et al., 2011, Ikeda et al., 2011, Peltzer et al., 2018, Tokunaga et al., 2011), we conjectured that HOIP could be unstable when recombinantly expressed in the absence of its interaction partners. To this end, we expressed HOIP (119.8 kDa), N-terminally His-tagged HOIL-1L (59.2 kDa), and N-terminally Strep(II)-tagged SHARPIN (43.0 kDa) in insect cells in order to co-purify the LUBAC holoenzyme by tandem affinity chromatography. Using this co-expression strategy, we were able to isolate three proteins of the expected molecular weights with no major contaminants as determined by SDS-PAGE followed by Coomassie staining (Figure 1B). Furthermore, we verified the identities of these proteins as the three LUBAC components by immunoblotting indicating the successful isolation of recombinant LUBAC (Figure 1C).

**Figure 1.**
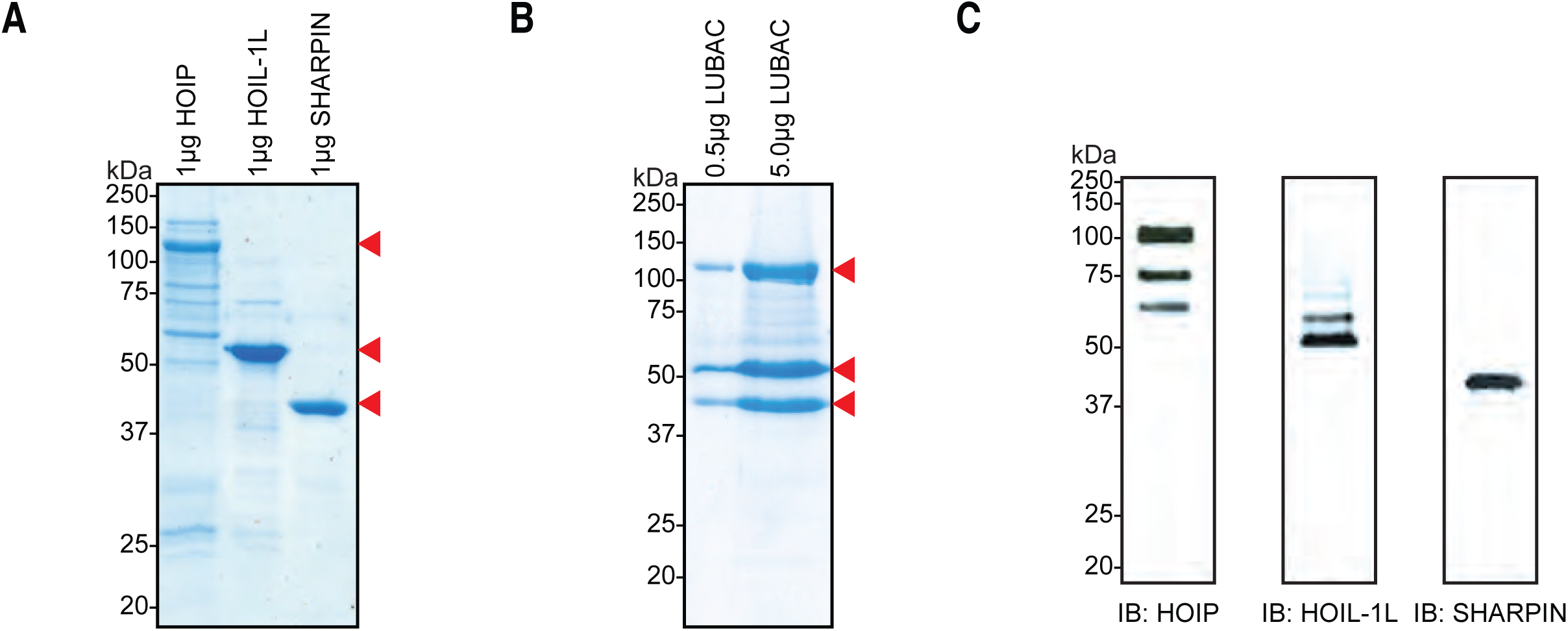
Co-expression and purification of LUBAC yields high quality protein. **A** SDS-PAGE analysis of individually purified LUBAC components. **B** SDS-PAGE analysis of co-expressed and purified LUBAC. **C** Immunoblot analysis of co-purified LUBAC.

To examine the stability of the purified complexes, we performed gel filtration chromatography (Figure EV1A). The elution profile of the complex contained two major peaks eluting at 0.942 ml (peak I) and 0.972 ml (peak II) as well as one minor peak eluting at 1.158 ml (peak III). All of these peaks eluted earlier than the 158 kDa molecular weight standard, which eluted at 1.246 ml (Figure EV1C). Given that the monomeric mass of the purified LUBAC complex is expected to be 222 kDa, the elution profile suggests that these peaks all correspond to assembled LUBAC in at least three populations of different oligomeric states. However, while peaks I and II contained all LUBAC components, peak III contained predominantly HOIL-1L and SHARPIN (Figure EV1A, Lower panel) indicating the presence of partially assembled complexes. To assess if this was a carryover form the purification or if the complex disassembles over time, we reapplied a fraction from peak II to the column (Figure EV1B). The elution profile from the tandem run contained almost exclusively assembled LUBAC (Figure EV1B, Lower panel) indicating that the complexes are not prone to dissociation after purification.

In order to screen the homogeneity of the sample, we imaged fractions from peak II by negative staining electron microscopy. Micrographs show a monodisperse distribution of particles of similar size, which could be sorted into 2D class averages showing a distinct elongated dumbbell structure thus verifying the homogeneity of the sample (Figure 2A and Figure EV2A). Furthermore, we were able to generate the first low-resolution 3D reconstruction of LUBAC from these particles (Figure 2B and Figure EV2B). In accordance with the class averages, the model displays an elongated asymmetric crescent structure with the majority of the mass concentrated at one end. The class averages match calculated projections of the generated model very well showing that the model is self-consistent. (Figure 2C and Figure EV3).

**Figure 2.**
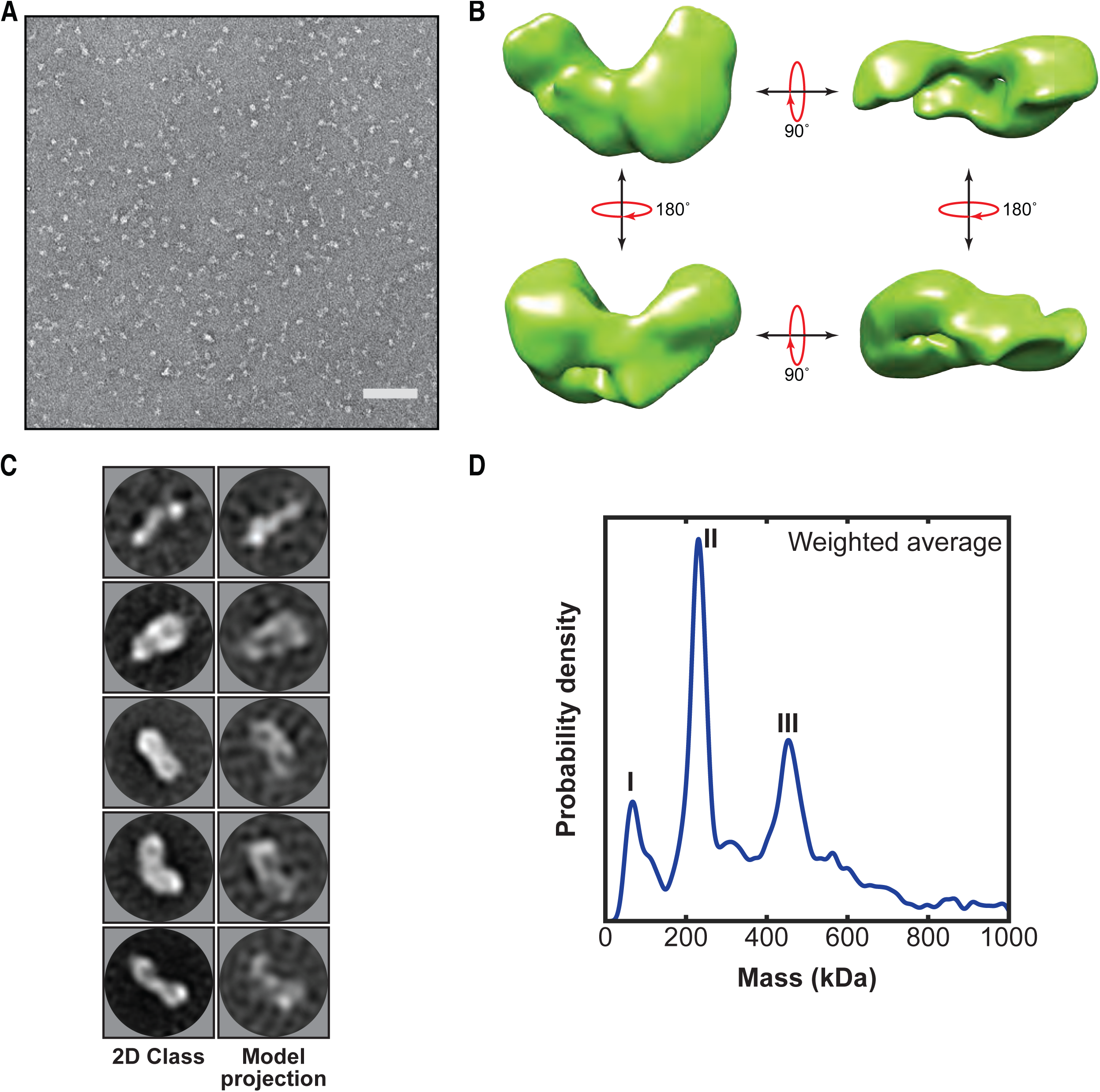
First low-resolution 3D map of LUBAC obtained by negative staining electron microscopy of the recombinant complex. **A** Representative negative stain transmission electron micrograph of recombinant LUBAC. Scale bar: 100 nm. **B**3D refined model of LUBAC obtained by single particle analysis of negative stained electron micrographs. **C** LUBAC 2D class averages matched to projections made from 3D refined map. **D** Mass photometry measurements of LUBAC indicate formation of a ternary complex with 1:1:1 stoichiometry that can form dimers.

Collectively, we established a purification protocol to obtain high purity and quality recombinant LUBAC that allowed us to generate the first low-resolution 3D reconstruction of the complex, from which we find the volume of structural envelope is consistent with a 230 kDa protein.

### The LUBAC complex exists as monomers and dimers with a 1:1:1 stoichiometry between HOIP, HOIL-1L, and SHARPIN

The precise stoichiometry and oligomerization state of the LUBAC complex have not been established although recent structural work has suggested a 1:1:1 stoichiometry between the three core components (Fujita et al., 2018). To determine the stoichiometry and oligomerization state of the LUBAC complex we used mass photometry (MP), a technique that enables accurate mass measurements based on the scattering of light by single macromolecules in solution (Sonn-Segev et al., 2019, Young et al., 2018) (Figure 2D and Figure EV4). MP measurements showed that the majority of species present in the samples had average masses of 231 kDa (Peak II), and 454 kDa (Peak III) (EV Table 1). An additional peak originating from a population with mass of less than 100 kDa was also measured (Peak I), which could arise from isolated HOIL-1L or SHARPIN present in the measured sample.

With respect to the expected mass of the ternary LUBAC complex the populations of peaks II and III nearly correspond with monomers (222 kDa) and dimers (444 kDa) of LUBAC with a 1:1:1 stoichiometry between HOIP, HOIL-1L, and SHARPIN.

### The RBR domains of HOIP and HOIL-1L are in close proximity

Understanding the precise mechanistic action of HOIL-1L and SHARPIN within LUBAC requires knowledge of structural and functional domain interactions between the three components. Current knowledge of the interaction domains of HOIP, HOIL-1L, and SHARPIN is shown in Figure 3A. Structural work of protein fragments has shown that HOIL-1L and SHARPIN interact with HOIP through their respective Ubiquitin-Like (UBL) domains, which bind cooperatively to the HOIP Ubiquitin-Associated (UBA) domain (Fujita et al., 2018, Liu et al., 2017, Yagi et al., 2012) (Figure 3A). By using truncation mutants, it has also been shown that the SHARPIN UBL domain interacts with the Npl Zinc Finger 2 (NZF2) domain of HOIP (Ikeda et al., 2011). More recently, structural work has revealed that SHARPIN and HOIL-1L interact with each other through their respective LUBAC-Tethering Motifs (LTMs) (Fujita et al., 2018). Otherwise, there is no further information available about the overall spatial arrangement of the domains of HOIP, HOIL-1L, and SHARPIN.

**Figure 3.**
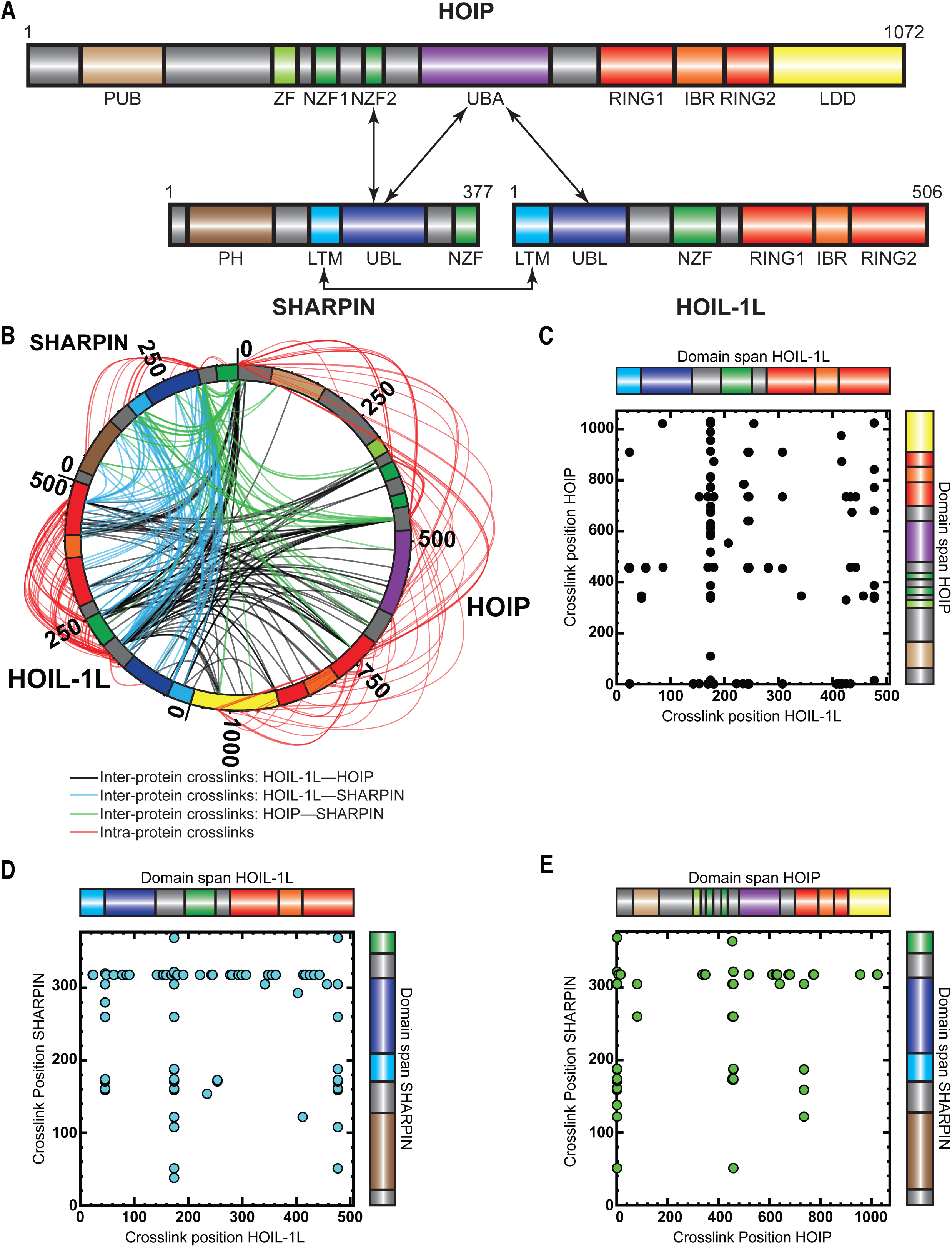
Cross-linking MS analysis shows proximity between the catalytic domains of HOIP and HOIL-1L. **A** Schematic representation of LUBAC components with their domains and known interactions. **B** Circos plot of inter-protein crosslinks formed between LUBAC components. **C** Detected inter-protein cross-links formed between HOIL-1L and HOIP. **D** Detected inter-protein cross-links formed between HOIL-1L and SHARPIN. **E** Detected inter-protein cross-links formed between HOIP and SHARPIN.

To obtain more detailed information about the spatial arrangement of LUBAC components, we performed cross-linking mass spectrometry (XL-MS) experiments. For this purpose, we used the amine-to-carboxyl-reactive cross linker ‘4-(4,6-Dimethoxy-1,3,5-triazin-2-yl)-4-methylmorpholinium tetrafluoroborate (DMTMM), a zero-length cross-linker (Leitner et al., 2014) revealing Lys and Asp/Glu contacts that are adjacent to each other. As shown in Figure 3B-E, we observed an extensive network of intra-protein and inter-protein crosslinks, providing a detailed picture of LUBAC complex assembly (Figure 3B-E, EV Tables 2 & 3). While some DMTMM cross-links were observed between LUBAC domains known to interact, such as between the HOIL-1L UBL and the SHARPIN UBL/LTM domains as well as between the HOIP UBA and HOIL-1L LTM domains, several cross-links pointed to new connections between the engaged proteins. For example, we observed crosslinks between the HOIL-1L RING1 and HOIP RING1/LDD domains, as well as between the HOIL-1L RING2 and the HOIP RING1/IBR/RING2/LDD domains. Additionally, HOIL-1L intra-protein crosslinks were formed between its NZF domain and its RING1/IBR/RING2 domains. In conclusion, the catalytic RBR domains of HOIP and HOIL1L seem to be close to each other, as well as the NZF and RBR regions of HOIL-1L. These data suggest that LUBAC may have a single catalytic centre containing the RBR domains of HOIP and HOIL-1L.

### Recombinant LUBAC assembles heterotypic ubiquitin chains containing linear and non-Lys linkages

To assess the ability of recombinant LUBAC to assemble linear ubiquitin chains, we performed *in vitro* ubiquitination assays. As expected, LUBAC generated linear ubiquitin chains in an ATP-dependent manner (Figure 4A). We also observed additional signals derived from co-purified LUBAC, which were absent from reactions containing the three individually purified and mixed LUBAC components (Figure 4B: red arrows).

**Figure 4.**
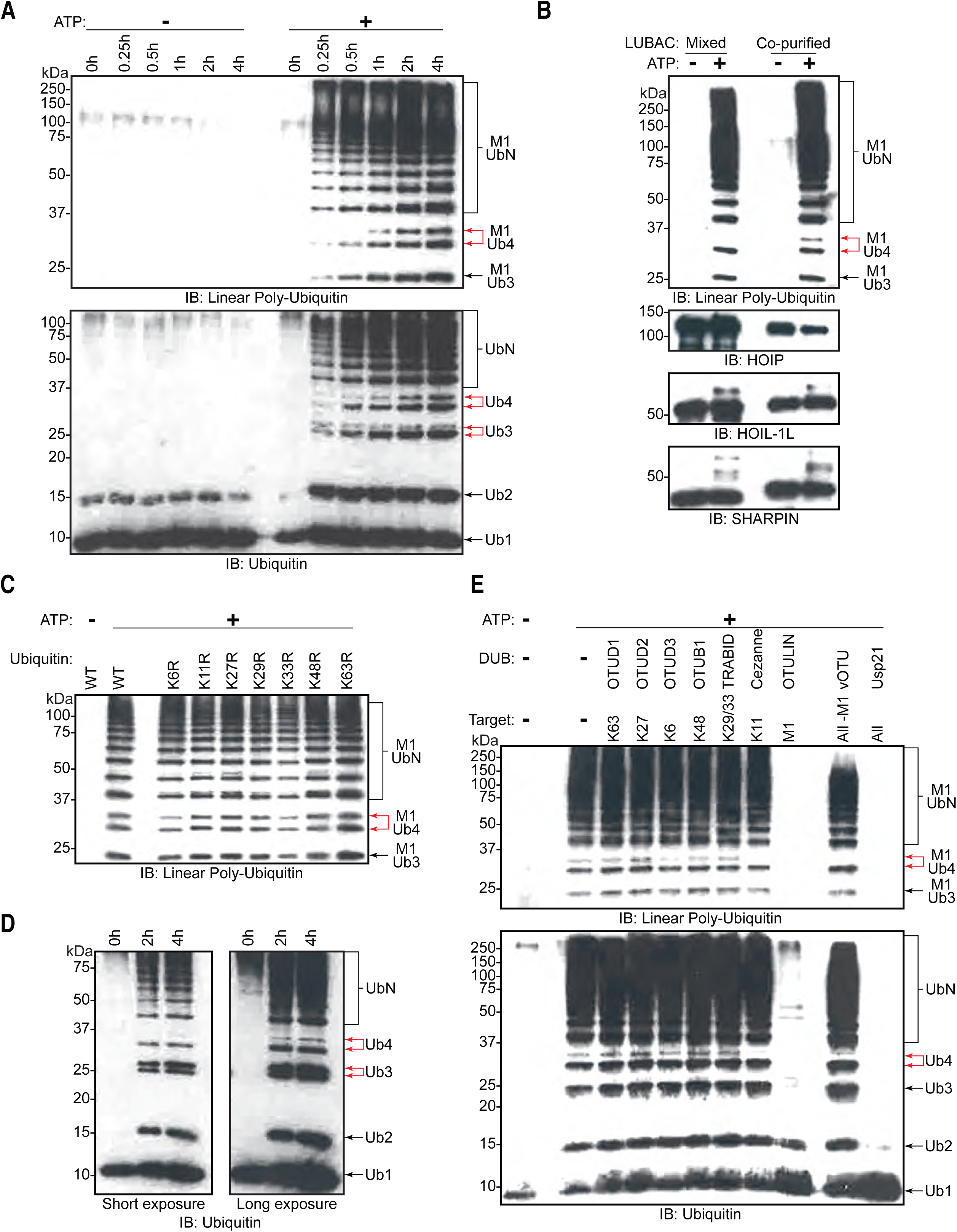
LUBAC assembles heterotypic poly-ubiquitin chains containing M1 and non-Lys linkages *in vitro*. **A** Time course of co-purified LUBAC *in vitro* ubiquitin chain assembly reaction. **B** Comparison of *in* vitro chain assembly between HOIP, HOIL-1L, and Sharpin mixed at 1:1:1 molar ratio versus co-purified LUBAC. **C** LUBAC *in vitro* chain assembly using different ubiquitin K to R mutants. **D** LUBAC *in vitro* chain assembly using K0 ubiquitin. **E** UbiCREST analysis of poly-ubiquitin chains assembled by LUBAC *in vitro*. All experiments were performed in triplicate representative results are shown.

Ubiquitin chains containing different linkages resolve at different apparent molecular masses by SDS-PAGE separation (Emmerich & Cohen, 2015). Therefore, we hypothesized that the two bands in each pair contained different ubiquitin linkages. To examine the presence of different Lys-linked bonds in the heterotypic chains, we carried out *in vitro* ubiquitination assays using different ubiquitin Lys to Arg (KR) mutants (Figure 4C). Heterotypic ubiquitin chain assembly was observed with all of the tested mutants, K6R, K11R, K27R, K29R, K33R, K48R and K63R. To rule out the possibility that mutation of a single Lys could be compensated by ubiquitination of a separate Lys residue, we also performed *in vitro* ubiquitination assay using ubiquitin lacking all Lys residues (K_0_) (Figure 4D). Despite reduced reaction efficiency, the K_0_ mutant could also be used by LUBAC to generate the heterotypic ubiquitin chains.

To further analyse the linkage types of ubiquitin chains, we performed UbiCRest (Hospenthal et al., 2015) on the ubiquitin chains generated by LUBAC (Figure 4E). UbiCrest using linkage-specific DUBs allows detection of specific ubiquitin chains by loss of signals when the linkage is targeted. Interestingly, the additional signals from ubiquitin chains assembled by LUBAC disappeared by treatment with Cezanne, a DUB specific for the Lys11-linkage (Bremm et al., 2010); or vOTU, a DUB targeting Lys-linkages (Akutsu et al., 2011) (Figure 4E; upper red arrow). The linear linkage-specific DUB OTULIN (Fiil et al., 2013) hydrolysed all linear ubiquitin bonds (Figure 4E: upper panel), corresponding to most of the ubiquitin signal, but left non-linear di- and tri-ubiquitin residues (Figure 4E: lower panel).

These data collectively indicate that recombinant LUBAC assembles heterotypic ubiquitin chains containing predominantly linear linkage with non-Lys-linked branches.

### LUBAC assembles heterotypic poly-ubiquitin chains containing linear and ester-linked bonds *in vitro* and in cells

A recent study showed that recombinant HOIL-1L can generate di-ubiquitin linked via an oxyester bond *in vitro*, and can also ubiquitinate substrates through oxyester bonds in cells (Kelsall et al., 2019). Therefore, we tested for the presence of oxyester bonds in the ubiquitin chains by checking their sensitivity to the nucleophile hydroxylamine (Figure 5A). Treatment with hydroxylamine resulted in the disappearance of the upper band of the linear tetra ubiquitin chain (Figure 5A: red arrow). This indicates that the chain branching is achieved by formation of an oxyester bond between ubiquitin moieties similar to the observation in the previous study (Kelsall et al., 2019).

**Figure 5.**
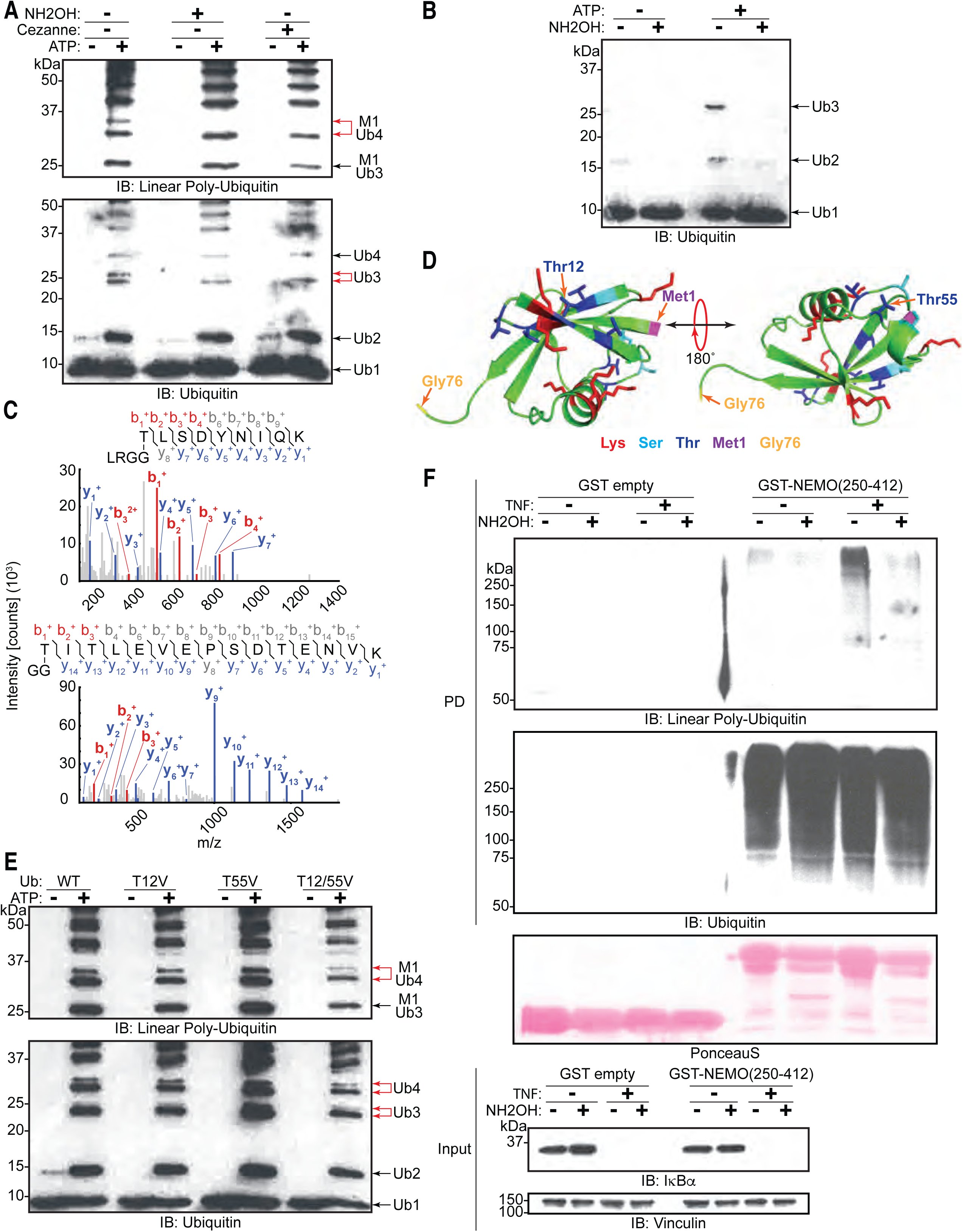
LUBAC assembles heterotypic poly-ubiquitin chains containing M1 bonds and ester bond linkages at T12 and T55. **A** Treatment of LUBAC-assembled heterotypic poly-ubiquitin chains with hydroxylamine. **B** Hydroxylamine treatment of ubiquitin polymers assembled by LUBAC using N-terminally blocked ubiquitin. **C** MS/MS spectra of ubiquitin polymerized at T55 (top) and T12 (bottom). Poly-ubiquitin chains assembled by LUBAC were separated by SDS-PAGE, bands were cut from the gel and subjected to mass spectrometry analysis. **D** Positions of Thr12 and Thr55 on structure of ubiquitin (PDB:1UBI). **E** Assembly of ubiquitin chains by LUBAC using different ubiquitin Thr to Val point mutants as substrates. **F** Hydroxylamine treatment of M1-containing poly-ubiquitin chains assembled in response to TNF in WT MEFs. MEFs were treated with TNF for 15 min and lysed, lysates were subjected to GST PD using GST or GST-NEMO(250-412), beads were treated with buffer or hydroxylamine for 30 min, bound ubiquitin species were then analysed by immunoblotting. All experiments were performed in triplicate representative results are shown.

To further ascertain whether LUBAC assembles ester-linked ubiquitin polymers, we carried out *in vitro* ubiquitination assays using N-terminally His_6_-tagged ubiquitin, which cannot be used to assemble linear ubiquitin chains (Kirisako et al., 2006). LUBAC inefficiently assembled di- and tri-ubiquitin chains with His-ubiquitin, which are sensitive to hydroxylamine treatment (Figure 5B). These results demonstrate that when the N-terminus of ubiquitin is not available, LUBAC can still assemble oxyester-linked poly-ubiquitin even in the absence of linear ubiquitin chains.

Ubiquitin contains seven Thr residues, three Ser residues, and one Tyr residue, which could theoretically act as sites for ester bond formation. Therefore, we aimed to identify the positions of the ester-linked branches using mass spectrometry by searching for GG dipeptides covalently attached through oxyester bonds to Ser, Thr, or Tyr. To this end, we analysed ubiquitin chains formed by LUBAC by LC-MS. MS/MS spectra of GG-conjugated dipeptides at residues Thr12 and Thr55 of ubiquitin were detected from these samples (Figure 5C). Structural analysis shows that these two residues are positioned at opposite sides of ubiquitin and neither of them is located in proximity to Met1 or the C-terminal Gly76 (Figure 5D; PDB:1UBI). This suggests that branches could potentially exist on both sites of a single ubiquitin molecule located at any position of a linear ubiquitin chain without creating any steric hindrances.

To further investigate if Thr12 and Thr55 are the sites of oxyester bond formation, we performed *in vitro* ubiquitination assays using ubiquitin mutated at Thr12 and/or Thr55 to Val (Figure 5E). Individual or concomitant mutation of Thr12 and Thr55 (Ub T_12_V, Ub T_55_V, or Ub T_12/55_V respectively) did not result in loss of chain branching. These results suggest that linear chain branching is not a strictly site-specific event.

We next probed the existence of these hybrid chains in cells. It has recently been reported that these chains are formed on IRAK1, IRAK2, and Myd88 in response to activation of TLR signalling (Kelsall et al., 2019). Given that LUBAC is involved in the TNF-signalling cascade, we examined the induction of these heterotypic ubiquitin chains in TNF-treated cells. Linear ubiquitin chains were enriched by GST pull-down using the NEMO-UBAN and Zinc finger domains (NEMO_250-412_) from total cell extracts of TNF stimulated MEFs and tested for hydroxylamine sensitivity (Figure 5F). TNF stimulation increased formation of linear ubiquitin chains detected by immunoblotting using an antibody specific for linear ubiquitin chains (Figure 5F: PD top panel lane 7). These chains proved sensitive to hydroxylamine treatment (Figure 5F: PD top panel lane 8). In contrast, the signals from the overall population of ubiquitin chains are unchanged by either TNF stimulation or hydroxylamine treatment (Figure 5F: PD bottom panel lanes 5-8). Together these results suggest that ubiquitin chains containing both linear linkages and hydroxylamine-sensitive ester bonds are produced in cells in response to TNF stimulation.

### HOIP assembles linear ubiquitin chains that are subsequently branched with ester bonds by HOIL-1L

We next proceeded to probe how HOIP and HOIL-1L generates these heterotypic ubiquitin chains. HOIP specifically assembles linear ubiquitin chains through the action of its RBR domains wherein Cys885 is the catalytic residue (Smit et al., 2012, Stieglitz et al., 2013); similarly HOIL-1L catalyses ester bond-directed ubiquitination via its RBR domain where Cys460 is the catalytic site (Kelsall et al., 2019, Smit et al., 2013). Therefore, we examined the possibility that HOIP assembles a linear ubiquitin chain, which is subsequently branched with ester bonds by HOIL-1L. To this end, we purified different LUBAC complexes containing catalytically inactive HOIP (HOIP C885A), HOIL-1L (HOIL-1L C460A), or both HOIP C885A and HOIL-1L C460A. Mutation of these residues did not impair the purification of LUBAC, and all variants of the complex could be isolated at similar yields and to similar degrees of purity (Figure EV5). In line with our hypothesis, LUBAC containing HOIL-1L C460A generated chains with linear linkages, yet the double band indicative of heterotypic chain assembly was absent (Figure 6A; lane 6 upper red arrow). These observations indicate that HOIL-1L is responsible for the formation of the ester-linked branches. Consistent with previous reports (Fuseya et al., 2020, Smit et al., 2013), HOIP assembled longer linear ubiquitin chains in the absence of HOIL-1L catalytic activity and did so more rapidly (Figure 6A: lanes 2 and 6). Conversely, LUBAC containing HOIP C885A was incapable of polymerizing ubiquitin altogether (Figure 6A; lanes 4 and 8) as expected. These results suggest that HOIL-1L catalytic activity disturbs linear ubiquitin chain formation but requires HOIP catalytic activity.

**Figure 6.**
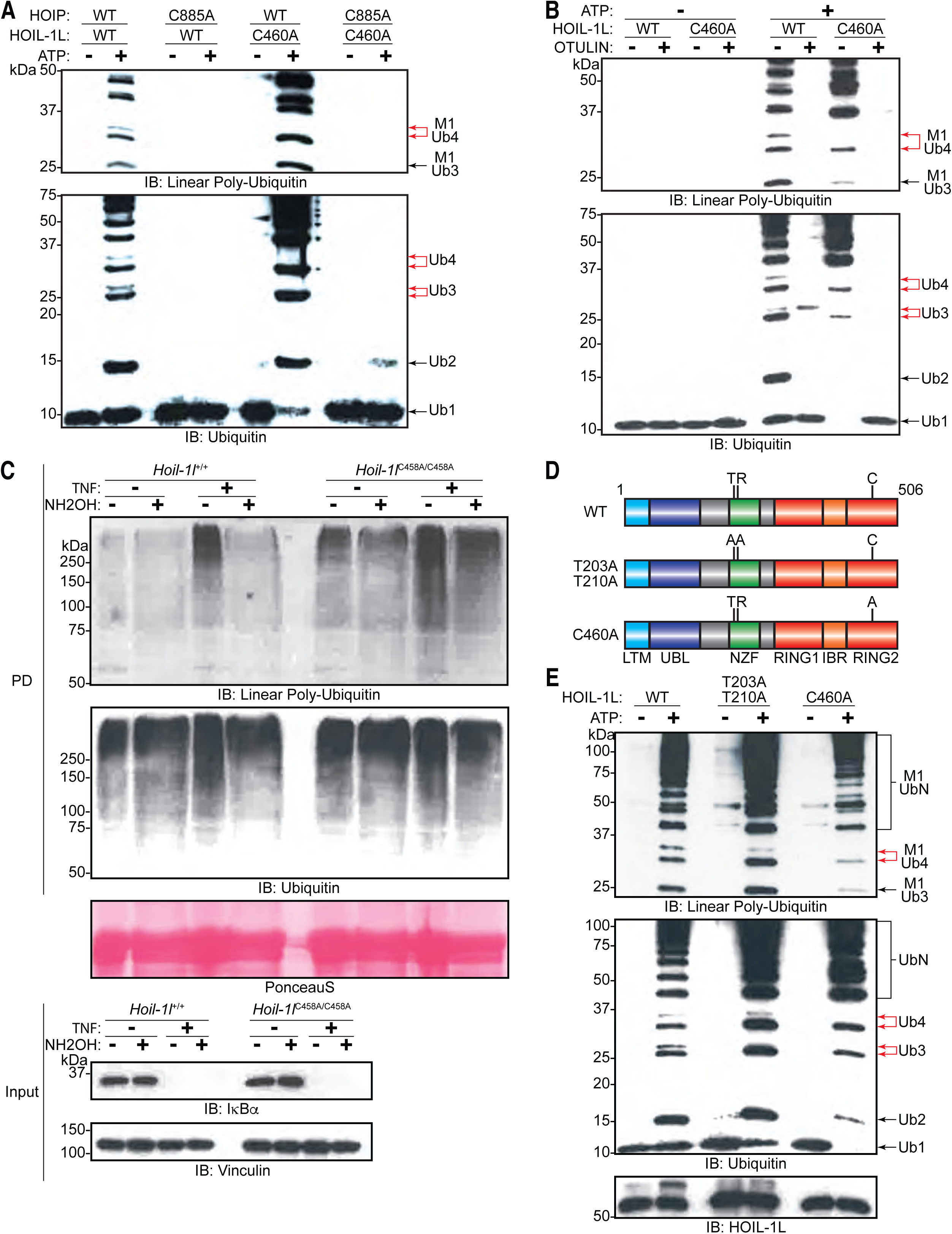
HOIL-1L generates ester linkages on heterotypic chains but requires HOIP catalytic activity to polymerize ubiquitin. **A** Comparison of LUBAC *in vitro* chain assembly by complexes containing different catalytically inert mutants of HOIP and HOIL-1L. **B** OTULIN restriction of poly-ubiquitin chains assembled by LUBAC containing WT or catalytically inert HOIL-1L. **C** Hydroxylamine treatment of M1-containing poly-ubiquitin chains assembled in response to TNF in WT and *Hoil-1l^C458A/C458A^* MEF cells. Cells were treated with TNF (50 ng/ml) for 15 min and lysed, lysates were subjected to GST PD using GST-NEMO(250-412), beads were treated with buffer or hydroxylamine for 30 min, bound ubiquitin species were then analysed by immunoblotting. **D** Schematic representations of different HOIL-1L mutants. **E** Comparison of LUBAC *in vitro* chain assembly by complexes containing different HOIL-1L mutants. All experiments were performed in triplicate representative results are shown.

The gel-migration pattern differed between ubiquitin chains assembled by LUBAC with HOIL-1L C460A and the WT complex (Figure 6A: lanes 2 and 6). Therefore, we compared the presence of non-linear bonds in the chains assembled by the different complexes by OTULIN. The restriction analyses showed that the ubiquitin chains assembled by LUBAC contained non-linear di- and tri-ubiquitin chains, whereas chains assembled by HOIL-1L C460A-containing LUBAC generated exclusively OTULIN-sensitive linear ubiquitin chains (Figure 6B). These results further support the claim that chain branching is dependent on the catalytic action of HOIL-1L while HOIP exclusively assembles linear ubiquitin chains.

### HOIL-1L catalytic activity is required for branched ubiquitin chain formation in cells

To further study the catalytic function of HOIL-1L in cells, we used cells derived from a HOIL-1L C458A knock-in mouse (*Hoil-1l^C458A/C458A^*) generated by CRISPR/Cas9 gene-editing technology (Figure EV 6A-C). The expression levels of the three LUBAC components are similar in mouse embryonic fibroblasts (MEFs) in *Hoil-1l^+/+^* and *Hoil-1l^C458A/C458A^* MEFs (Figure EV6D). In agreement with numerous studies from the past, two species of HOIL-1L were detected in *Hoil-1l^+/+^* MEFs by immunoblotting; the slower migrating species originating from auto-ubiquitinated HOIL-1L was absent in *Hoil-1l^C458A/C458A^* MEFs.

To determine if HOIL-1L is the responsible ligase for linear ubiquitin chain branching during TNF signalling, TNF-treated *Hoil-1l^C458A/C458A^* MEF lysates were subjected to GST pull-down with GST-NEMO_250-412_ followed by hydroxylamine treatment (Figure 6C). Similarly to WT MEFs, TNF stimulation led to an increase in the levels of linear ubiquitin chains in *Hoil-1l^C458A/C458A^* MEFs (Figure 6C: PD top panel lanes 3 and 7). However, the enriched ubiquitin signals detected by immunoblotting revealed a higher basal level of linear ubiquitin chains in *Hoil-1l^C458A/C458A^* MEFs compared to those in *Hoil-1l^+/+^* MEFs (Figure 6C: PD top panel lanes 1 and 5). Moreover, unlike in *Hoil-1l^+/+^* MEFs, linear ubiquitin chains in *Hoil-1l^C458A/C458A^* MEFs were not sensitive to degradation by hydroxylamine treatment (Figure 6C: PD bottom panel). However, there was no obvious difference when blotting for general ubiquitin between *Hoil-1l^+/+^* and *Hoil-1l^C458A/C458A^* MEFs regardless of TNF stimulation or chain hydroxylamine treatment (Figure 6C: PD bottom panel). These results show that ester-linked branching of linear ubiquitin chains formed during TNF signalling in MEFs is dependent on the catalytic activity of HOIL-1L.

### HOIL-1L Npl4 Zinc Finger (NZF) is involved in the formation of branching ubiquitin chains *in vitro*

Since the catalytic action of HOIP precedes that of HOIL-1L and the assembly of linear ubiquitin chains precedes the appearance of the branches, we hypothesized that HOIL-1L interacts with a linear ubiquitin chain as a substrate for branching via its linear ubiquitin chain-specific binding domain NZF (Sato et al., 2011). To test this possibility, we purified LUBAC containing a mutant in which critical residues for linear ubiquitin chain recognition are mutated (HOIL-1L T203A,R210A) (Figure 6D). In agreement with our hypothesis, chain branching activity by LUBAC was partially impaired by the HOIL-1L T203A,R210A mutant when compared to the WT or HOIL-1L C460A (Figure 6E). Interestingly, LUBAC with HOIL-1L T203A,R210A assembled ubiquitin chains more efficiently than WT LUBAC but less efficiently than HOIL-1L C460 (Figure 6E).

These data collectively show that HOIP assembles linear ubiquitin chains, which are subsequently branched by HOIL-1L in a process involving its NZF domain and which requires the catalytic activity of HOIP.

In summary, we identified that LUBAC assembles heterotypic linear/ester-linked poly-ubiquitin chains *in vitro* and in cells in response to TNF stimulation. We also show that these chains are synthesized through the concerted action of HOIP and HOIL-1L (Figure 7). These chains may contribute in modulating the speed and/or efficiency of linear ubiquitin chain synthesis by LUBAC.

**Figure 7.**
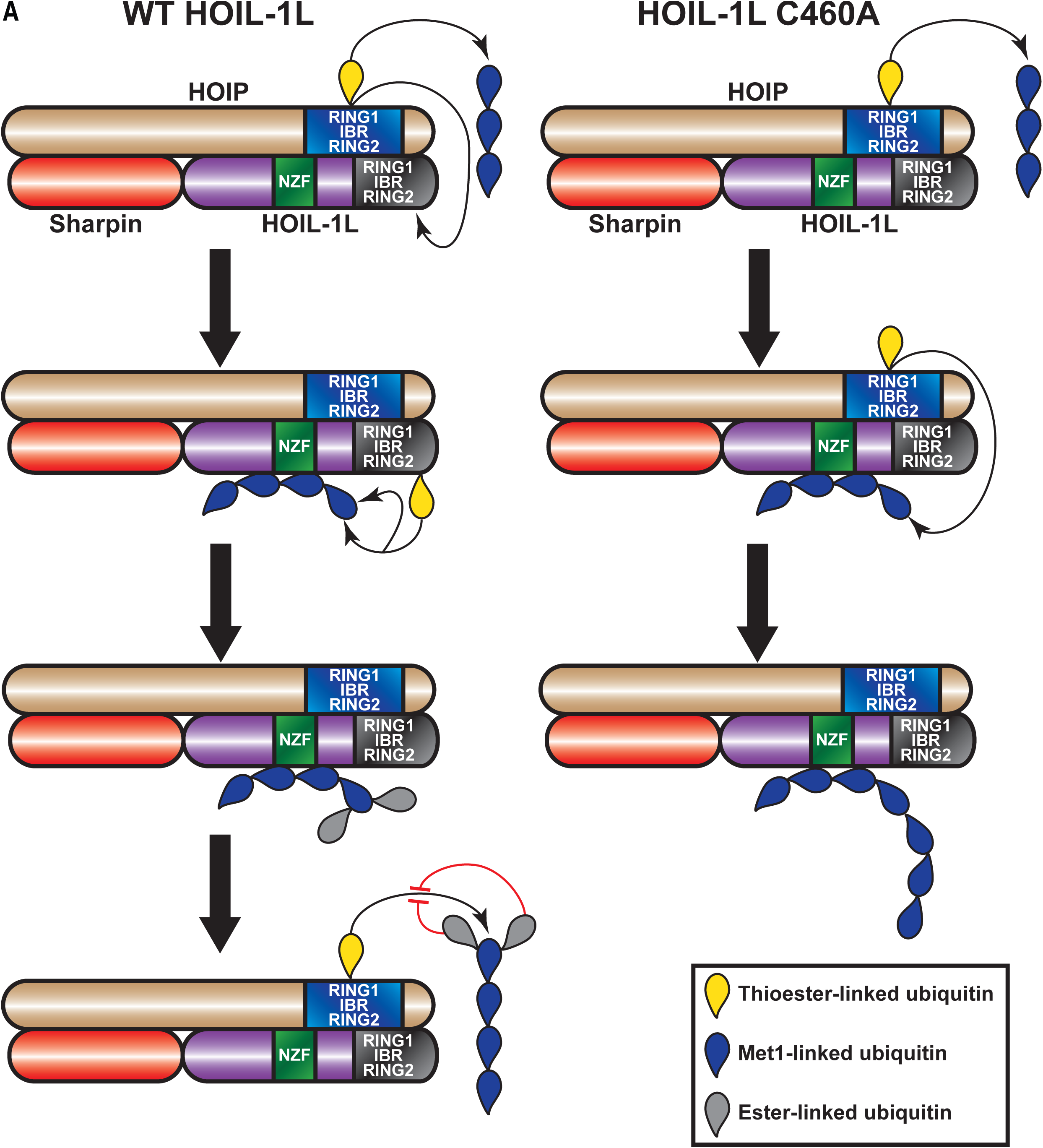
Proposed model for concerted action between HOIP and HOIL-1L in heterotypic chain assembly. HOIP and HOIL-1L assemble heterotypic chains through a Cys-relay mechanism. HOIP forms a thioester bond to ubiquitin, which can be either transferred to a thioester bond on HOIL-1L or added to a nascent M1-linked chain. HOIL-1L subsequently binds the M1-linked chain through its NZF domain and branches it with ester-linkages. The resulting heterotypic poly-ubiquitin chains contain predominantly M1 linkages with ester-linked branches.

## Discussion

We present here the first 3D reconstruction of the ternary LUBAC complex. We cannot determine the exact positions of HOIP, HOIL-1L, and SHARPIN in the map at the current resolution; however, our map is the first structure encompassing the LUBAC holoenzyme. We also made other novel structural observations that LUBAC exists as monomers and dimers of a ternary complex with 1:1:1 stoichiometry between HOIP, HOIL-1L, and SHARPIN. This is in agreement with observations made in recent structural work (Fujita et al., 2018). We also present new data addressing the question of LUBAC oligomerization. Early gel filtration analysis of cell-derived LUBAC suggested formation of a large or oligomeric complex with a molecular mass of over 600 kDa (Kirisako et al., 2006). However, our MP measurements using recombinant LUBAC show that this is not the case *in vitro*. This could be due to LUBAC in cells interacting with cellular components such as SPATA2 and CYLD (Elliott et al., 2016, Kupka et al., 2016, Schlicher et al., 2016, Wagner et al., 2016). This discrepancy would also be accounted for by a particle of non-globular structure, which would elute earlier than expected from a gel filtration column (Erickson, 2009, Siegel & Monty, 1966). Accordingly, the 3D map we obtained from negative stain electron micrographs show a particle of elongated structure.

We propose a catalytic Cys relay mechanism in LUBAC: HOIP receives ubiquitin from the E2 and uses this ubiquitin to assemble linear chains or transfers it HOIL-1L, which then branches the chains assisted by its NZF domain (Figure 7). One recent example of such a mechanism is the E3 ligase MYCBP2, which also conjugates the ubiquitin to Thr residues through an ester bond (Pao et al., 2018). A relay mechanism would require spatial proximity between the catalytic sites of the two ligases. Our XL-MS data show that this is indeed the case based on crosslinking between the RBR domains of HOIP and HOIL-1L. We also detected proximity between the HOIL-1L NZF and the RBR domains of both HOIP and HOIL-1L, which could contribute to the branching of the linearly linked ubiquitin dimer. Collectively, these observations suggest that the LUBAC complex has a single catalytic centre containing the RBR domains of HOIP and HOIL-1L.

HOIP and HOIL-1L have distinct catalytic behaviour. HOIP exclusively targets Met1 of a ubiquitin moiety for polymerization (Swatek et al., 2019); in contrast, HOIL-1L seems to have more flexibility with target sites in forming ester bonds. We identified Thr12 and Thr55 in ubiquitin as target sites by mass spectrometry; yet individual and concomitant mutation of these residues did not abolish heterotypic chain assembly suggesting existence of secondary sites. Indeed, in ubiquitin, Thr12 is structurally located in the vicinity of Thr7, Thr9, and Thr14, whereas Thr55 is located near Thr22, Ser20, and Ser57. A recent study identified Thr12, Ser20 and Thr22, but not Thr55, as target sites by isolated HOIL-1L (Kelsall et al., 2019). These differences may depend on HOIL-1L as an isolated protein, or HOIL-1L as a part of LUBAC where HOIP and SHARPIN contribute to site selection.

We speculate that HOIP and SHARPIN contribute to the catalytic action of HOIL-1L based on the observed loss of the HOIL-1L auto-ubiquitination signal in cells derived from SHARPIN-deficient mice (Gerlach et al., 2011, Ikeda et al., 2011, Tokunaga et al., 2011) and HOIP knockout mice (Peltzer et al., 2014). This effect is identical in cells derived from HOIL-1L C458S mice (Kelsall et al., 2019), as well as HOIL-1L C458A mice from this study. Investigating the precise mechanism how the non-catalytic LUBAC component SHARPIN contributes to the HOIL-1L catalytic activity may shed light on its role in the complex.

DUBs targeting ester linkages remain to be identified. Our data show that a fragment of Cezanne, a DUB specific for Lys11 linkages (Bremm et al., 2010, Mevissen et al., 2016), can cleave the ester-linked branches *in vitro*. Cezanne is a known negative regulator of NF-κB (Enesa et al., 2008, Evans et al., 2003, Evans et al., 2001, Luong le et al., 2013). It is therefore tempting to speculate that it has esterase activity and targets the LUBAC-assembled chains during its counteraction of NF-κB activation. However, further work will be necessary to identify potential esterase activity in DUBs including Cezanne.

Understanding the biological functions of the heterotypic chains is a very important aspect. A recent study using HOIL-1L ΔRBR mice and HOIL-1L knockin cell of ligase inactive mutant showed that HOIL-1L ligase activity regulates TNF-induced signaling and apoptosis (Fuseya et al., 2020). In line with their observations, we also observed that HOIL-1L C458 mutant as a part of LUBAC increased the linear ubiquitination signal in comparison to the wild type control. They also showed that RIPK1, a known substrate of LUBAC, in the TNF complex is linearly ubiquitinated in at increased levels. These data indicate that the HOIL-1L catalytic activity negatively regulates ubiquitination of the substrates in cells, which also supports our observations *in vitro*. In cells, they could not detect oxyester-linkage on HOIL-1L but on HOIP in wild type cells, which is abolished in HOIL-1L catalytic inactive cells. Since they detect remaining HOIL-1L mono-ubiquitination signal even when all the Lys residues in HOIL-1L are mutated to Arg, it is most probably that there are heterogeneous population of ubiquitination sites in HOIL-1L. Furthermore, this could be due to that oxyester bond being biochemically less stable and being a minor fraction in cells. Since HOIL-1L inactive mutant is no longer ubiquitinated *in vitro*, upregulated ubiquitination of HOIL-1L mutant in cells (Fuseya et al., 2020) may depend on interacting partners existing in cells. To further understand the roles of these ester-linked branches in biology, it will be important to dissect biochemical characteristics such as their interaction mode with known linear ubiquitin chain specific binding domains (Fennell et al., 2018), existence of DUBs, and other possible ligases that catalyse their formation in the future studies.

## Materials and Methods

### Plasmids

pGEX-6p1-*Hs*OTULIN, pGEX-6p1-*Hs*HOIP, pGEX-6p1-*Hs*HOIL-1L and pGEX-6p1-*Hs*SHARPIN (Fennell et al., 2019), as well as pGEX-4t1-mNEMO_250-412_ (Wagner et al., 2008) have been previously reported. pGEX-6p1-*Hs*UbcH7 was kindly provided by Katrin Rittinger (Francis Crick Institute, UK). pOPIN-K-*Hs*OTUD1_287-481_ (Mevissen et al., 2013), pOPIN-K-*Hs*OTUD2 (Mevissen et al., 2013), pOPIN-K-*Hs*OTUD3_52-209_ (Mevissen et al., 2013), pOPIN-K-*Hs*Cezanne_53-446_ (Mevissen et al., 2013), pOPIN-K-*Hs*TRABID_245-697_ (Licchesi et al., 2011), pOPIN-K-*Hs*OTUB1 (Mevissen et al., 2013), pOPIN-K-vOTU_1-183_ (Akutsu et al., 2011), and pOPIN-S-*Hs*Usp21_196-565_ (Ye et al., 2011) were kind gifts from David Komander. ORF of Ubiquitin T_12_V, T_55_V, and T_12/55_V mutants were generated by PCR and inserted twice into a pETDuet-1 vector by isothermal assembly. pKL-der-*Hs*LUBAC was assembled by inserting *Hs*HOIP, His_6_-*Hs*HOIL-1L, and Strep(II)-*Hs*Sharpin coding sequences into a pKL-derived vector using BsaI. pKL-der-*Hs*LUBAC-HOIP(C_885_A), pKL-der-*Hs*LUBAC-HOIL-1L(C_460_A), pKL-der-*Hs*LUBAC-HOIP(C_885_A)-HOIL-1L(C_460_A), and pKL-der-*Hs*LUBAC-HOIL-1L(T_203_A/R_210_A) were generated by a standard protocol of site-directed mutagenesis.

### Antibodies

The following antibodies were used in this study: anti-HOIL-1L (clone 2E2; Merck MABC576), anti-HOIP (Merck SAB2102031), anti-IκBα (44D4; Cell Signalling Technology 4812), anti-Ser32/36-phospho-IκBα (5A5; Cell Signaling Technology 9246), anti-SHARPIN (NOVUS Biologicals NBP2-04116), anti-ubiquitin (P4D1; Santa Cruz Biotechnology sc-8017), anti-vinculin (Merck V9131). All antibodies were diluted in TBS 5% (w/v) BSA, 0.05% (v/v) Triton according to the manufacturer’s recommended dilutions.

### Generation of anti-linear ubiquitin monoclonal antibody

Mouse monoclonal anti-linear ubiquitin chain antibody (clone LUB4), was generated by immunizing five-week-old male ICR mice (Charles River Laboratory, Yokohama, Japan) with a neoepitope peptide (LRLRGGMQIFVK) derived from the linear ubiquitin chain, which comprises a ubiquitin C-terminal sequence (amino acids 71-76) and a ubiquitin N-terminal sequence (amino acids 1-6) conjugated to OVA. Cells isolated from the popliteal and inguinal lymph nodes were fused with a mouse myeloma cell line, PAI. Supernatants of the growing hybridomas were tested by direct enzyme-linked immunosorbent assay (ELISA) and western blot analysis. Specificity of the antibody clone was validated by immunoblotting (Figure EV7).

### Immunoblotting

Protein samples were mixed with SDS-sample buffer containing 5% β-mercaptoethanol and denatured at 96°C for 5 minutes. Proteins were separated by SDS-PAGE and transferred to nitrocellulose membranes (GE Healthcare 10600019 or 10600001). Appropriate transfer of proteins was verified by staining of membranes with the Ponceau S solution (Roth 5938.1). Membranes were blocked with TBS 5% BSA (w/v), 0.05% Triton (v/v) and blotted at 4°C overnight with indicated primary antibodies. Membranes were subsequently blotted with anti-mouse-IgG-HRP conjugate (Bio-Rad 1706516) or anti-rabbit-HRP conjugate (Agilent Dako P044801-2) and visualized with Western Blotting Luminol Reagent (Santa Cruz Biotechnology sc-2048) using Amersham Hyperfilm MP (GE Healthcare) chemiluminescence films.

### Purification of recombinant LUBAC from insect cells

The transfer vector carrying the HsHOIP, His6-HsHOIL-1L, and Strep(II)-HsSHARPIN coding sequences was cloned using the GoldenBac cloning system (Neuhold et al., 2020) and transposed into the bacmid backbone carried by DH10EmBacY cells (Geneva Biotech). The bacmid was then used to generate a V0 virus stock in Sf9 cells, which was amplified once to give a V1 virus stock used to infect 1 l cultures of Sf9 cells (Expression Systems). Cells were grown in ESF 921 Insect Cell Culture Medium Protein Free (expression systems 96-001-01) at 27°C and infected at a density of 2×10^6^ cells/ml. Cells were harvested 72 hrs after growth arrest and resuspended in 100 mM HEPES, 100 mM NaCl, 100 mM KCl, 100 mM Arg, 10 mM MgSO_4_, 20 mM Imidazole, pH 8.0 supplemented with 1 tablet of cOmplete Mini EDTA-free protease inhibitor cocktail (Roth). Cells were lysed by 4 passes through a Constant Systems Cell Disruptor at 1.4 kBar and then supplemented with 100 mM Benzoase and 10 mM PMSF. Lysates were cleared by centrifugation at 48,284 g for 45 min at 4°C and loaded into a HisTrap FF cartridge (GE Healthcare). Proteins were eluted with 100 mM HEPES, 100 mM NaCl, 100 mM KCl, 50 mM Arginine, 500 mM imidazole, pH 8.0 and loaded into a Streptactin Superflow cartridge (IBA Lifesciences 2-1238-001). Cartridge was washed with 100 mM HEPES, 100 mM NaCl, 100 mM KCl, pH 8.0 and proteins were eluted with 100 mM HEPES, 100 mM NaCl, 100 mM KCl, 5 mM D-desthiobiotin pH 8.0. Complexes were concentrated using a Centriprep (Merck 4311) centrifugal filter with a 50 kDa cut-off then flash frozen in liquid nitrogen and stored at −80°C.

### Protein purification from *E. coli*

Proteins were expressed in *E. coli* BL21(DE3) cells (Agilent) for 16 hrs at 18°C, 25°C, or 30°C. Cell pellets were harvested and lysed by sonication with a Branson 450 Digital Sonifier (Branson) by pulsing for 1.5 min with 1 sec pulses and 2 sec pauses at amplitude of 60%. Cells expressing GST-HOIP, GST-HOIL-1L, GST-SHARPIN, or GST-UbcH7 were lysed in 100 mM HEPES, 500 mM NaCl, pH 7.5 supplemented with 1 tablet of cOmplete Mini EDTA-free protease inhibitor cocktail. Cells expressing ubiquitin mutants were lysed in 50 mM ammonium acetate pH 4.5 supplemented with 1 tablet of cOmplete Mini EDTA-free protease inhibitor cocktail. Cells expressing His_6_-SUMO-Usp21_196-565_ were lysed in 50 mM Tris, 200mM NaCl, pH 8.5 supplemented with 1 tablet of cOmplete Mini EDTA-free protease inhibitor cocktail. Cells expressing GST-OTUD1_287-481_, GST-OTUD2, GST-OTUD3_52-_ 209, GST-Cezanne_53-446_, GST-TRABID_245-697_, GST-OTUB1, GST-OTULIN, and GST-vOTU_1-183_ were lysed in 50 mM Tris 200 mM NaCl, 5 mM DTT, pH 8.5 supplemented with 1 tablet of cOmplete Mini EDTA-free protease inhibitor cocktail. Cell lysates were supplemented with 500 U Recombinant DNAse (Merck 4716728001) and 100 mM PMSF. For purification of HOIP, HOIL-1L, SHARPIN, and UbcH7 lysates were loaded into a GSTrap HP cartridge (GE Healthcare); once loaded GST-PreScission Protease (GE Healthcare) was injected into the column and incubated overnight at 4°C. UbcH7, HOIP, HOIL-1L, and Sharpin were eluted and further purified over a Superdex 75 16/600 pd or a Superdex 200 16/600 pd column (GE Healthcare) equilibrated in 50 mM HEPES, 150 mM NaCl, pH 7.5. For ubiquitin purification, lysates were loaded into a ResourceS (GE Healthcare) cartridge and eluted in a gradient of 0 to 500 mM NaCl in 50 mM ammonium acetate, pH 4.5. The protein was further purified over a Superdex 75 16/600 pd column (GE Healthcare) equilibrated in 50 mM HEPES, 150 mM NaCl, pH 7.5. OTUD1_287-481_, OTUD2, OTUD3_52-209_, Cezanne_53-446_, TRABID_245-697_, OTUB1, OTULIN, and vOTU_1-183_ were purified as described in (Hospenthal et al., 2015), in brief: lysates were loaded into a GSTrap HP cartridge (GE Healthcare) and treated overnight with GST-PreScission Protease (GE Healthcare). Eluted proteins were further purified over a Superdex 75 16/600 pd column (GE Healthcare) equilibrated in 50 mM Tris, 200 mM NaCl 5 mM DTT, pH 8.5. For purification of Usp21_196-565_ lysate was loaded ino a HisTrap FF cartridge (GE Healthcare) and eluted with 50 mM Tris, 200 mM NaCl, 500 mM Imidazole, pH 8.5. Imidazole was removed by buffer exchange using a Vivaspin centrifugal filter (Sartorius) with a 10 kDa cut-off, eluate was treated with SENP2 (R&D Systems E-710-050) overnight and protein was further purified over a Superdex 75 16/600 pd column (GE Healthcare) equilibrated in 50 mM Tris, 200mM NaCl, 5 mM DTT, pH 8.5. Proteins were concentrated using Vivaspin centrifugal filters (Sartorius) of appropriate cut-offs, flash frozen in liquid nitrogen and stored at −80°C. For GST and GST-NEMO_250-412_, they were expressed in BL21 (DE3) cells for 16 hrs at 25°C (Wagner et al., 2008). Cells were harvested and lysed by sonication in 50 mM Tris, 100 mM EDTA, 50 mM EGTA, 150 mM NaCl, pH 7.5 supplemented with 1 tablet of cOmplete Mini EDTA-free protease inhibitor cocktail, lysates were incubated overnight at 4°C with 400 μl of Glutathione Sepharose 4B per l of expression culture (GE Healthcare GE17-0756-01) with gentle rotation. Beads immobilized samples were washed in 50 mM Tris, 100 mM EDTA, 150 mM NaCl, 0.5% Triton, pH 7.5 and resuspended in 50 mM Tris, pH 7.5 (Einarson & Orlinick, 2002).

### GST pull-down assays and NH_2_OH treatment

MEFs were lysed in lysis buffer containing 50 mM HEPES, 150 mM NaCl, 1 mM EDTA, 1 mM EGTA, 1% (v/v) Triton X-100, 10% (v/v) Glycerol, 25 mM NAF, 10 μM ZnCl_2_, 10 mM NEM, 1 mM PMSF, 5 mM Na_3_VO_4_, pH 7.4 supplemented with 1 tablet of cOmplete Mini EDTA-free protease inhibitor cocktail. Lysates were cleared by centrifugation at 21,130 g for 15 min at 4°C; Equal amounts of GST or GST-NEMO_250-412_, as determined by SDS-PAGE, were added to lysates and samples were incubated overnight at 4°C with gentle rocking. Beads washed with PBS then buffer containing 100 mM HEPES, 100 mM NaCl, 100 mM KCl, pH 8.0 then resuspended in a buffer containing 100 mM HEPES, 100 mM NaCl, 100 mM KCl, pH 9.0 ±1.2 M NH_2_OH. Reactions were incubated 30 min at 37°C in a Thermomixer comfort (Eppendorf, Germany) thermomixer; reactions were stopped with SDS buffer and incubated at 37°C then subjected to SDS-PAGE and immunoblotting with the indicated antibodies.

### Negative staining

Recombinant LUBAC was run through a Superdex S200 Increase 3.2/300 column equilibrated in 100 mM HEPES, 100 mM NaCl, 100 mM KCl, pH 8.0. Fractions containing the peaks for recombinant LUBAC were collected for staining on 400-mesh Cu/Pd grids (Agar Scientific G2440PF) coated with a 3 nm thick continuous carbon support film. For staining grids were glow discharged for 1 min at 200 mA and 10^−1^ mBar in a BAL-TEC SCD005 sputter coater (BAL-TEC, Liechtenstein), samples were pipetted onto the grid and incubated for 1 min before blotting. Grids were then stained with 2% (w/v) uranyl acetate pH 4 (Merck) for 1 min, blotted dry, and left to air-dry for 10 min at room temperature and pressure before being stored in a desiccator.

### Electron microscopy

Grids were screened on a FEI Morgani 268D transmission electron microscope (Thermo Fisher Scientific, The Netherlands) operated at 80 kV using a 300 μm condenser lens aperture and 50 μm objective lens aperture; images were recorded on an 11-megapixel Morada CCD camera (Olympus-SIS, Germany). For data acquisition grids were imaged on a FEI Tecnai G2 20 (Thermo Fisher Scientific, The Netherlands) operated at 200 kV using a 200 μm condenser lens aperture and 100 μm objective lens aperture; imaged on a FEI Eagle 4k HS CCD camera (Thermo Fisher Scientific, The Netherlands) with a pixel size of 1.85 Å/pixel.

### Image processing of negative stain data

The negative stain data was processed in relion 3.1 (Scheres, 2012a, Scheres, 2012b, Zivanov et al., 2018) unless otherwise stated. CTF parameters were determined using CTFFIND4 (Rohou & Grigorieff, 2015). Micrographs that suffered from drift were excluded from the further analysis. 600 particles were picked manually and classified in 2D to yield templates for autopicking. Autopicking yielded 637,000 particles that were extracted in a box of 180 pixels but cropped in Fourier space to 90 pixels. The dataset was split in 6 subsets that were subjected to 2D classification. Class averages showing distinct particle features were kept and their particles were combined for a final round of 2D classification yielding 163,000 remaining particles. These particles were subjected to initial model generation yielding 3 models that are very similar in shape. The most isotropic model was used for 3D classification once again yielding similar models. All steps were performed without CTF correction to avoid artifacts and over-fitting. Back projections of the final map were generated to validate the self-consistency of the model.

### Crosslinking mass spectrometry

For crosslinking preparations 100 μg of recombinant LUBAC were cross-linked in 100 mM HEPES, 100 mM NaCl, 100 mM KCl, pH 8.0 with 3.75 mM DMTMM (Merck 74104) for 1 hr at RT. The reaction cross-linker was removed using a Zeba Micro Spin desalting column (Thermo Fisher Scientific 89883). Samples were dried at 45°C in a vacuum centrifuge connected to a cooling trap (Heto Lab Equipment) and then resuspended in 8 M Urea. Proteins were subjected to reduction with 2.5 mM TCEP and subsequently alkylated with 5 mM iodoacetamide for 30 min at RT after which samples were diluted 1:7 (v/v) with 50 mM ammonium bicarbonate. Samples were digested with 2 μg Trypsin (Promega V5280) for 20 hrs at 37°C, samples were then digested for a further 3 hrs with 2 μg Chymotrypsin (Roche 11418467001) at 25°C. Digest was quenched with 0.4% (v/v) trifluoroacetic acid (TFA) and samples were loaded on a Sep-Pak C18 cartridge (Waters WAT054955) equilibrated with 5% (v/v) acetonitrile, 0.1% (v/v) formic acid (FA); samples were eluted with 50% (v/v) acetonitrile, 0.1% (v/v) FA. Cross-linked peptides were enriched in a Superdex 30 Increase 3.2/300 column (GE Healthcare) equilibrated with 30% (v/v) acetonitrile, 0.1% (v/v) TFA. Fractions containing cross-linked peptides were evaporated to dryness and resuspended in 5% (v/v) acetonitrile, 0.1% (v/v) TFA. Cross-linked peptides were separated on an UltiMate 3000 RSLC nano HPLC system (Thermo Fisher Scientific) coupled to an Orbitrap Fusion Lumos Tribid mass spectrometer (Thermo Fisher Scientific) equipped with a Proxeon nanospray source (Thermo Fisher Scientific). Peptides were loaded onto an Acclaim PepMap 100 C18 HPLC column (Thermo Fisher Scientific 160454) in a 0.1% TFA mobile phase. Peptides were eluted into an Acclaim PepMap 100 C18 HPLC column (Thermo Fisher Scientific 164739) in a binary gradient between mobile phase A (99.9/0.1% v/v water/FA) and mobile phase B (19.92/80/0.08% v/v/v water/acetonitrile/FA). The gradient was run from 2-45% mobile phase B over 3 hrs and then increased to 90% B over 5 min.

Measurement was performed in data-dependent mode with 3 sec cycle time at a resolution of 60,000 in the m/z range of 350-1500. Precursor ions with a charge state of +3 to +7 were fragmented by HCD with collision energy of 29% and fragments were recorded with a resolution of 45000 and precursor isolation width of 1.0 m/z. A dynamic exclusion time of 30 sec was used. Fragment spectra peak lists were generated with MSConvert v3.0.9974 (Chambers et al., 2012) with a peak-picking filter. Crosslink identification was done using XiSearch v1.6.742 (Giese et al., 2016) with 6 ppm MS^1^-accuracy, 20 ppm MS^2^-accuracy, DMTMM-cross linking was set with asymmetric reaction specificity for lysine and arginine to glutamate and aspartate; carbamidomethylation of cysteine was a fixed modification, oxidation of methionine was a variable modification, trypsin/chymotrypsin digest was set with up to 4 missed cleavages, and all other variables were left at default settings. Identified cross-links were filtered to 10% FDR at the link level with XiFDR v1.1.27 (Fischer & Rappsilber, 2017), links with FDR<5% were visualized with xVis (Grimm et al., 2015).

### *In vitro* ubiquitination assay

Reaction mixtures were prepared containing Ube1, UbcH7, and either LUBAC or HOIP, HOIL-1L, and SHARPIN with all proteins at 0.338 μM final concentration in a buffer containing 50 mM HEPES, 150 mM NaCl, 0.5 mM MgSO_4_, pH 8.0 (described in Ikeda et al. 2011). Reactions were started by addition of 59.0 μM ubiquitin and 2 mM ATP then incubated at 37°C in a Mastercycler nexus (Eppendorf, Germany) thermocycler for 2 hrs unless otherwise stated. Reaction was stopped by addition of SDS buffer and boiling at 95°C, proteins were resolved by SDS-PAGE, and analysed by immunoblotting with the indicated antibodies. Recombinant human ubiquitin, His_6_-ubiquitin, and ubiquitin KR mutants (K_6_R, K_11_R, K_27_R, K_29_R, K_33_R, K_48_R, K_63_R, K_0_) were purchased from Boston Biochem.

### NH2OH treatment of *in vitro* assembled poly-ubiquitin chains

LUBAC *in vitro* reactions were mixed with 1.2 M NH_2_OH in 40 mM HEPES, 40 mM NaCl, 40 mM KCl, pH 9.0 and reactions were incubated at 37°C in a Thermomixer comfort (Eppendorf, Germany) thermomixer for 2 hrs. Reactions were stopped by addition of SDS buffer and incubated at 37°C then subjected to SDS-PAGE and immunoblotting with the indicated antibodies.

### Ubiquitin Chain Restriction (UbiCRest) assay

Protocol was adapted from (Hospenthal et al., 2015); in brief: deubiquitinases (DUBs) were diluted to 0.8 μM (OTUD1_287-481_, OTUD2, OTUD3_52-209_, Cezanne_53-446_, TRABID_245-697_, OTUB1, and OTULIN) or 10 μM (vOTU_1-183_, Usp21_196-565_) in 25 mM Tris, 150 mM NaCl, 10 mM DTT, pH 7.5 and incubated at RT for 10 min. One LUBAC *in vitro* chain assembly reaction was divided into aliquots and mixed at a 1:1 (v/v) ratio with each DUB in 50 mM Tris, 50 mM NaCl, 5 mM DTT, pH 7.5 and reactions were incubated at 37°C in a Thermomixer comfort (Eppendorf, Germany) thermomixer for 2 hrs. Reactions were stopped as described above, and analysed by immunoblotting with the indicated antibodies.

### Mass spectrometry

LUBAC *in vitro* chain assembly reactions were resolved on 4-15% Mini-PROTEAN TGX gels (Bio-Rad 4561083) and stained with Instant Blue (Merck ISB1L). Fragments were excised from the gel and slices were washed sequentially with 100 mM ammonium bicarbonate (ABC) then 50% (v/v) acetonitrile 50 mM ABC at 57°C for 30 min. Gel slices were shrunk in 100% acetonitrile and reduced with 1 mg/ml DTT in 100 mM ABC. Slices were then alkylated with MMTS in 100 mM ABC at RT for 30 min. Gel slices were subjected to tryptic digest (Promega V5280) overnight at 37°C. Peptides were extracted from gel slices by sonication in 5% (v/v) formic acid (FA). Peptides were loaded into a PepMap C18 (5 mm × 300 μm ID, 5 μm particles, 100 Å pore size; Thermo Fisher Scientific) trap column on an UltiMate 3000 RSLC nano HPLC system (Thermo Fisher Scientific, Amsterdam, Netherlands) coupled to a Q Exactive HF-X mass spectrometer (Thermo Fisher Scientific, Bremen, Germany) equipped with a Proxeon nanospray source (Thermo Fisher Scientific, Odense, Denmark) using a solution of 0.1% (v/v) TFA as the mobile phase. Samples were loaded at a flow rate of 25 μl/min for 10 min and then eluted into an analytical C18 (500 mm × 75 μm ID, 2 μm, 100 Å; Thermo Fisher Scientific, Amsterdam, Netherlands) column in a binary gradient between mobile phase A (99.9/0.1% v/v water/FA) and mobile phase B (19.92/80/0.08% v/v/v water/acetonitrile/FA). The gradient was run from 98%/2% A/B to 35%/65% A/B over 1 hour; the gradient was then adjusted to 5%/95% A/B over 5 minutes and held for a further 5 minutes before returning to 98%/2% A/B. The Q Exactive HF-X mass spectrometer was operated in data-dependent mode, using a full scan (m/z range 380-1500, nominal resolution of 60,000, target value 1_E_^6^) followed by MS/MS scans of the 10 most abundant ions. MS/MS spectra were acquired using normalized collision energy of 28, isolation width of 1.0 m/z, resolution of 30,000 and target value was set to 1_E_^5^. Precursor ions selected for fragmentation were put on a dynamic exclusion list for 20 sec (excluding charge states 1, 7, 8, >8). The minimum AGC target was set to 5 ^3^ and intensity threshold was calculated to be 4.8_E_^4^; the peptide match feature was set to preferred and the exclude isotopes feature was enabled. For peptide identification, the generated .raw files were loaded into Proteome Discoverer (version 2.3.0.523, Thermo Fisher Scientific). All hereby created MS/MS spectra were searched using MSAmanda v2.0.0. 12368 (Dorfer et al., 2014). The .raw files were searched against the *Spodoptera frugiperda* database (23,492 sequences; 12,713,298 residues) and the swissprot-ecoli database (4,418 sequences; 1,386,900 residues) to generate a .fasta file; search parameters used were: peptide mass tolerance ±5 ppm, fragment mass tolerance 15 ppm, maximum of 2 missed cleavages; the result was filtered to 1% FDR on protein level using Percolator algorithm integrated in Thermo Proteome Discoverer. The sub-database was generated for further processing. The .raw files were then searched using the generated .fasta file using the following search parameters: β-methylthiolation on cysteine was set as a fixed modification; oxidation of methionine, deamidation of asparagine and glutamine, acetylation of lysine, phosphorylation of serine, threonine and tyrosine; methylation of lysine and arginine, di-methylation of lysine and arginine, tri-methylation of lysine, biotinylation of lysine, carbamylation of lysine, ubiquitination of lysine, serine, threonine, and tyrosine were set as variable modifications. Monoisotopic masses were searched within unrestricted protein masses for tryptic enzymatic specificity. The peptide mass tolerance was set to ±5 ppm, fragment mass tolerance to ±15 ppm, maximum number of missed cleavages was set to 2, results were filtered to 1% FDR on protein level using Percolator algorithm (Kall et al., 2007) as integrated in Proteome Discoverer. The localization of the post-translational modification sites within the peptides was performed with the tool ptmRS, based on the tool phosphoRS (Taus et al., 2011). Peptide areas were quantified using in-house-developed tool apQuant (Doblmann et al., 2019).

### Generation of C57BL/6J *Hoil-1l*^C458A/C458A^ mice and genotyping

C57BL/6J *Hoil-1l^C458A/C458A^* mice were generated as described elsewhere (Fennell et al, BioRxiv). The gRNA targeting sequence was designed using the online tool ChopChop (http://chopchop.cbu.uib.no), two separate gRNA sequences were selected (gRNA1 and gRNA63). The gRNA sequences were inserted to pX330-U6-Chimeric_BB-CBh-hSpCas9 (a gift from Feng Zhang, Addgene plasmid # 42230) (Cong et al., 2013) as annealed oligonucleotides (EV Table 4) using BbsI. A T7-gRNA product was amplified by PCR and used for *in vitro* transcription using the MEGAshortscript T7 kit (Invitrogen AM1345). The *in vitro* transcribed gRNA was purified with the MEGAclear kit (Invitrogen AM1908). A single-stranded donor template oligonucleotide (ssOligo) was designed containing the C458A mutation, silent mutation of the PAM site and a silent Hpy188III restriction site (5’ CGGATTGTGGTCCAGAAGAAAGACGGCGCTGACTGGATTCGCTGTACAGTCTGCCACACTGAGATC 3’). Superovulation was induced in 3-5 week old female C57BL/6J donor mice by treatment with 5IU of pregnant mare’s serum gonadotropin (Hölzel Diagnostika OPPA01037) and 5IU of human chorionic gonadotropin (Intervet, GesmbH); females were mated and zygotes were isolated in M2 media (Merck Milipore MR-015P-D). Zygotes were cultured in KSOM medium (Cosmo Bio Co., Ltd RB074) and cytosolically injected with 100 ng/μl Cas9 mRNA (Merck CAS9MRNA-1EA), 50 ng/μl gRNA, and 200 ng/μl ssOligo; injected zygotes were transferred to pseudo-pregnant females. Correct knock in was confirmed in founder mice by Sanger sequencing. Routine genotyping of mice was done by digestion with Hpy188III (NEB R0622) and analysis on a 2% agarose gel.

### Mouse husbandry

C57BL/6J *Hoil-1l^C458A/C458A^* mice were bred and kept at the IMBA/IMP animal house. All procedures with animals were carried according to institutional, Austrian, and European guidelines.

### Isolation and immortalization of MEFs

Mouse Embryonic Fibroblast (MEF) isolation and immortalization methods are described elsewhere (Fennell et al., 2019). Briefly, primary MEFs were isolated at E13.5 following standard protocols. Pregnant females were sacrificed, and the uterus was extracted. Isolated tissue from the embryo was digested with trypsin (Thermo Fisher 25300054) for 5 min at 37°C and digest was quenched with DMEM containing 10% (v/v) FCS. Cells were collected by centrifugation and cultured at 37°C, 5% CO2 in Dulbecco’s Modified Eagle’s Medium (DMEM; Sigma D5648) supplemented with 10% (v/v) FCS, 0.2 U/ml penicillin, 0.2 μg/ml streptomycin (Merck P0781), and 2 mM L-glutamine (Gibco 25030-024). MEFs were immortalized with SV40 large T-antigen by transfection using GeneJuice transfection reagent (Merck Millipore 70967).

### TNF stimulation of MEFs

1.6×106 MEFs were seeded in a 10 cm cell culture dish and grown overnight, serum-starved overnight in DMEM supplemented with 0.2% (v/v) FCS, 0.2 U/ml penicillin, 0.2 μg/ml streptomycin, and 2 mM L-glutamine. The following day cells were stimulated for 15 min with 50 ng/ml mouse TNF (Peprotech).

## Acknowledgements

We thank Dr Gijs Versteeg (Max F. Perutz Laboratories, Austria) and Katrin Rittinger (The Crick Insisute, UK) for critical discussion and feedback on the work and manuscript. We thank Olga Olszanska, Steven Dupart and Lilian Fennell (IMBA, Austria) for their technical support. We also thank Richard Imre, Elisabeth Roitinger, and Otto Hudecz for mass spectrometry data analysis, as well as Ines Steinmacher and Susanne Opravil for technical support (the Protein Chemistry Facility, IMP/IMBA core facility, Austria). Samples were prepared and data was recorded at the EM Facility of the Vienna BioCenter Core Facilities GmbH (VBCF, Austria). We thank the IMP/IMBA core facilities, including Transgenic Service, Comparative Medicine, and Molecular Biology Service for their technical support. Baculovirus production and insect cell culture was performed by the Protein Technologies Facility (VBCF, Austria). Research at the Ikeda Lab is supported by JSPS KAKENHI (Grant Number JP 18K19959), the ERC Consolidator Grant (LUbi, 614711), FWF Stand-Alone Grant (P 2550 8), and the Austrian Academy of Sciences. Research at the Haselbach Lab is supported by Boehringer Ingelheim. Research at the Clausen Lab is supported by the FFG Headquarter Grant 852936 and Boehringer Ingelheim. Research at the Kukura Lab is supported by the ERC Starting Grant PHOTOMASS 819593.

## Author contributions

A.R.C. planned and performed experiments of LUBAC purification, *in vitro* biochemical assays, and cellular assays. A.R.C. and A.V. prepared samples and carried out data analysis for XL-MS experiments. C.G.D. and K.S. contributed to knock in mouse generation and MEF isolation, A.R.C. and L.D. purified recombinant proteins, Z.O.-N. made measurements for XL-MS experiments, A.S.-S. carried out mass photometry measurements and data analysis in the lab of P.K.. S.S. generated an anti-linear ubiquitin chain antibody clone. A.R.C. and D.H. generated and processed negative staining EM data sets. K.M. contributed to MS experiments. A.R.C., A.V., D.H., and F.I. made figures and wrote the manuscript. D.H., F.I., and T.C. planned, organized, and coordinated the project, and supervised author students.

## References

Akutsu M, Ye Y, Virdee S, Chin JW, Komander D (2011) Molecular basis for ubiquitin and ISG15 cross-reactivity in viral ovarian tumor domains. Proc Natl Acad Sci U S A 108: 2228–33

Bhogaraju S, Kalayil S, Liu Y, Bonn F, Colby T, Matic I, Dikic I (2016) Phosphoribosylation of Ubiquitin Promotes Serine Ubiquitination and Impairs Conventional Ubiquitination. Cell 167: 1636–1649 e13

Bremm A, Freund SMV, Komander D (2010) Lys11-linked ubiquitin chains adopt compact conformations and are preferentially hydrolyzed by the deubiquitinase Cezanne. Nature Structural & Molecular Biology 17: 939–947

Chambers MC, Maclean B, Burke R, Amodei D, Ruderman DL, Neumann S, Gatto L, Fischer B, Pratt B, Egertson J, Hoff K, Kessner D, Tasman N, Shulman N, Frewen B, Baker TA, Brusniak M-Y, Paulse C, Creasy D, Flashner L et al. (2012) A cross-platform toolkit for mass spectrometry and proteomics. Nature Biotechnology 30: 918–920

Cong L, Ran FA, Cox D, Lin S, Barretto R, Habib N, Hsu PD, Wu X, Jiang W, Marraffini LA, Zhang F (2013) Multiplex genome engineering using CRISPR/Cas systems. Science 339: 819–23

Doblmann J, Dusberger F, Imre R, Hudecz O, Stanek F, Mechtler K, Durnberger G (2019) apQuant: Accurate Label-Free Quantification by Quality Filtering. J Proteome Res 18: 535–541

Dorfer V, Pichler P, Stranzl T, Stadlmann J, Taus T, Winkler S, Mechtler K (2014) MS Amanda, a universal identification algorithm optimized for high accuracy tandem mass spectra. J Proteome Res 13: 3679–84

Einarson MB, Orlinick JR (2002) Identification of protein-protein interactions with glutathione-S-transferase fusion proteins. Protein-protein interactions: a molecular cloning manual Cold Spring Harbor Laboratory Press, Cold Spring Harbor, NY: 37–52

Elliott PR, Leske D, Hrdinka M, Bagola K, Fiil BK, McLaughlin SH, Wagstaff J, Volkmar N, Christianson JC, Kessler BM, Freund SM, Komander D, Gyrd-Hansen M (2016) SPATA2 Links CYLD to LUBAC, Activates CYLD, and Controls LUBAC Signaling. Mol Cell 63: 990–1005

Emmerich CH, Cohen P (2015) Optimising methods for the preservation, capture and identification of ubiquitin chains and ubiquitylated proteins by immunoblotting. Biochemical and Biophysical Research Communications 466: 1–14

Enesa K, Zakkar M, Chaudhury H, Luong le A, Rawlinson L, Mason JC, Haskard DO, Dean JL, Evans PC (2008) NF-kappaB suppression by the deubiquitinating enzyme Cezanne: a novel negative feedback loop in pro-inflammatory signaling. J Biol Chem 283: 7036–45

Erickson HP (2009) Size and shape of protein molecules at the nanometer level determined by sedimentation, gel filtration, and electron microscopy. Biol Proced Online 11: 32–51

Evans PC, Smith TS, Lai MJ, Williams MG, Burke DF, Heyninck K, Kreike MM, Beyaert R, Blundell TL, Kilshaw PJ (2003) A novel type of deubiquitinating enzyme. J Biol Chem 278: 23180–6

Evans PC, Taylor ER, Coadwell J, Heyninck K, Beyaert R, Kilshaw PJ (2001) Isolation and characterization of two novel A20-like proteins. Biochem J 357: 617–23

Fennell LM, Deszcz L, Schleiffer A, Mechtler K, Kavirayani A, Ikeda F (2019) Site-specific ubiquitination of the E3 ligase HOIP regulates cell death and immune signaling. bioRxiv: 742544

Fennell LM, Rahighi S, Ikeda F (2018) Linear ubiquitin chain-binding domains. FEBS J 285: 2746–2761

Fiil BK, Damgaard RB, Wagner SA, Keusekotten K, Fritsch M, Bekker-Jensen S, Mailand N, Choudhary C, Komander D, Gyrd-Hansen M (2013) OTULIN restricts Met1-linked ubiquitination to control innate immune signaling. Mol Cell 50: 818–830

Fischer L, Rappsilber J (2017) Quirks of Error Estimation in Cross-Linking/Mass Spectrometry. Anal Chem 89: 3829–3833

Fujita H, Tokunaga A, Shimizu S, Whiting AL, Aguilar-Alonso F, Takagi K, Walinda E, Sasaki Y, Shimokawa T, Mizushima T, Ohki I, Ariyoshi M, Tochio H, Bernal F, Shirakawa M, Iwai K (2018) Cooperative Domain Formation by Homologous Motifs in HOIL-1L and SHARPIN Plays A Crucial Role in LUBAC Stabilization. Cell Reports 23: 1192–1204

Fuseya Y, Fujita H, Kim M, Ohtake F, Nishide A, Sasaki K, Saeki Y, Tanaka K, Takahashi R, Iwai K (2020) The HOIL-1L ligase modulates immune signalling and cell death via monoubiquitination of LUBAC. Nat Cell Biol

Gerlach B, Cordier SM, Schmukle AC, Emmerich CH, Rieser E, Haas TL, Webb AI, Rickard JA, Anderton H, Wong WW, Nachbur U, Gangoda L, Warnken U, Purcell AW, Silke J, Walczak H (2011) Linear ubiquitination prevents inflammation and regulates immune signalling. Nature 471: 591–6

Giese SH, Fischer L, Rappsilber J (2016) A Study into the Collision-induced Dissociation (CID) Behavior of Cross-Linked Peptides. Mol Cell Proteomics 15: 1094–104

Grimm M, Zimniak T, Kahraman A, Herzog F (2015) xVis: a web server for the schematic visualization and interpretation of crosslink-derived spatial restraints. Nucleic Acids Res 43: W362–9

Hospenthal MK, Mevissen TET, Komander D (2015) Deubiquitinase-based analysis of ubiquitin chain architecture using Ubiquitin Chain Restriction (UbiCRest). Nat Protoc 10: 349–361

Ikeda F (2015) Linear ubiquitination signals in adaptive immune responses. Immunological reviews 266: 222–36

Ikeda F, Deribe YL, Skanland SS, Stieglitz B, Grabbe C, Franz-Wachtel M, van Wijk SJ, Goswami P, Nagy V, Terzic J, Tokunaga F, Androulidaki A, Nakagawa T, Pasparakis M, Iwai K, Sundberg JP, Schaefer L, Rittinger K, Macek B, Dikic I (2011) SHARPIN forms a linear ubiquitin ligase complex regulating NF-kappaB activity and apoptosis. Nature 471: 637–41

Iwai K, Tokunaga F (2009) Linear polyubiquitination: a new regulator of NF-kappaB activation. EMBO Rep 10: 706–13

Kall L, Canterbury JD, Weston J, Noble WS, MacCoss MJ (2007) Semi-supervised learning for peptide identification from shotgun proteomics datasets. Nat Methods 4: 923–5

Kelsall IR, Zhang J, Knebel A, Arthur JSC, Cohen P (2019) The E3 ligase HOIL-1 catalyses ester bond formation between ubiquitin and components of the Myddosome in mammalian cells. Proceedings of the National Academy of Sciences 116: 13293–13298

Kirisako T, Kamei K, Murata S, Kato M, Fukumoto H, Kanie M, Sano S, Tokunaga F, Tanaka K, Iwai K (2006) A ubiquitin ligase complex assembles linear polyubiquitin chains. EMBO J 25: 4877–87

Kupka S, De Miguel D, Draber P, Martino L, Surinova S, Rittinger K, Walczak H (2016) SPATA2-Mediated Binding of CYLD to HOIP Enables CYLD Recruitment to Signaling Complexes. Cell Reports 16: 2271–2280

Leitner A, Joachimiak LA, Unverdorben P, Walzthoeni T, Frydman J, Förster F, Aebersold R (2014) Chemical cross-linking/mass spectrometry targeting acidic residues in proteins and protein complexes. Proceedings of the National Academy of Sciences 111: 9455–9460

Licchesi JD, Mieszczanek J, Mevissen TE, Rutherford TJ, Akutsu M, Virdee S, El Oualid F, Chin JW, Ovaa H, Bienz M, Komander D (2011) An ankyrin-repeat ubiquitin-binding domain determines TRABID’s specificity for atypical ubiquitin chains. Nat Struct Mol Biol 19: 62–71

Liu J, Wang Y, Gong Y, Fu T, Hu S, Zhou Z, Pan L (2017) Structural Insights into SHARPIN-Mediated Activation of HOIP for the Linear Ubiquitin Chain Assembly. Cell Rep 21: 27–36

Luong le A, Fragiadaki M, Smith J, Boyle J, Lutz J, Dean JL, Harten S, Ashcroft M, Walmsley SR, Haskard DO, Maxwell PH, Walczak H, Pusey C, Evans PC (2013) Cezanne regulates inflammatory responses to hypoxia in endothelial cells by targeting TRAF6 for deubiquitination. Circ Res 112: 1583–91

Mevissen TE, Hospenthal MK, Geurink PP, Elliott PR, Akutsu M, Arnaudo N, Ekkebus R, Kulathu Y, Wauer T, El Oualid F, Freund SM, Ovaa H, Komander D (2013) OTU deubiquitinases reveal mechanisms of linkage specificity and enable ubiquitin chain restriction analysis. Cell 154: 169–84

Mevissen TET, Kulathu Y, Mulder MPC, Geurink PP, Maslen SL, Gersch M, Elliott PR, Burke JE, van Tol BDM, Akutsu M, Oualid FE, Kawasaki M, Freund SMV, Ovaa H, Komander D (2016) Molecular basis of Lys11-polyubiquitin specificity in the deubiquitinase Cezanne. Nature 538: 402–405

Neuhold J, Radakovics K, Lehner A, Weissmann F, Garcia MQ, Romero MC, Berrow NS, Stolt-Bergner P (2020) GoldenBac: a simple, highly efficient, and widely applicable system for construction of multi-gene expression vectors for use with the baculovirus expression vector system. BMC Biotechnol 20: 26

Noad J, von der Malsburg A, Pathe C, Michel MA, Komander D, Randow F (2017) LUBAC-synthesized linear ubiquitin chains restrict cytosol-invading bacteria by activating autophagy and NF-kappaB. Nat Microbiol 2: 17063

Pao KC, Wood NT, Knebel A, Rafie K, Stanley M, Mabbitt PD, Sundaramoorthy R, Hofmann K, van Aalten DMF, Virdee S (2018) Activity-based E3 ligase profiling uncovers an E3 ligase with esterification activity. Nature 556: 381–385

Peltzer N, Darding M, Montinaro A, Draber P, Draberova H, Kupka S, Rieser E, Fisher A, Hutchinson C, Taraborrelli L, Hartwig T, Lafont E, Haas TL, Shimizu Y, Boiers C, Sarr A, Rickard J, Alvarez-Diaz S, Ashworth MT, Beal A et al. (2018) LUBAC is essential for embryogenesis by preventing cell death and enabling haematopoiesis. Nature 557: 112–117

Peltzer N, Rieser E, Taraborrelli L, Draber P, Darding M, Pernaute B, Shimizu Y, Sarr A, Draberova H, Montinaro A, Martinez-Barbera JP, Silke J, Rodriguez TA, Walczak H (2014) HOIP deficiency causes embryonic lethality by aberrant TNFR1-mediated endothelial cell death. Cell Rep 9: 153–165

Qiu J, Sheedlo MJ, Yu K, Tan Y, Nakayasu ES, Das C, Liu X, Luo ZQ (2016) Ubiquitination independent of E1 and E2 enzymes by bacterial effectors. Nature 533: 120–4

Rahighi S, Ikeda F, Kawasaki M, Akutsu M, Suzuki N, Kato R, Kensche T, Uejima T, Bloor S, Komander D, Randow F, Wakatsuki S, Dikic I (2009) Specific Recognition of Linear Ubiquitin Chains by NEMO Is Important for NF-κB Activation. Cell 136: 1098–1109

Rittinger K, Ikeda F (2017) Linear ubiquitin chains: enzymes, mechanisms and biology. Open biology 7

Rivkin E, Almeida SM, Ceccarelli DF, Juang YC, MacLean TA, Srikumar T, Huang H, Dunham WH, Fukumura R, Xie G, Gondo Y, Raught B, Gingras AC, Sicheri F, Cordes SP (2013) The linear ubiquitin-specific deubiquitinase gumby regulates angiogenesis. Nature 498: 318–24

Rohou A, Grigorieff N (2015) CTFFIND4: Fast and accurate defocus estimation from electron micrographs. J Struct Biol 192: 216–21

Sato Y, Fujita H, Yoshikawa A, Yamashita M, Yamagata A, Kaiser SE, Iwai K, Fukai S (2011) Specific recognition of linear ubiquitin chains by the Npl4 zinc finger (NZF) domain of the HOIL-1L subunit of the linear ubiquitin chain assembly complex. Proc Natl Acad Sci U S A 108: 20520–5

Scheres SH (2012a) A Bayesian view on cryo-EM structure determination. J Mol Biol 415: 406–18

Scheres SH (2012b) RELION: implementation of a Bayesian approach to cryo-EM structure determination. J Struct Biol 180: 519–30

Schlicher L, Wissler M, Preiss F, Brauns-Schubert P, Jakob C, Dumit V, Borner C, Dengjel J, Maurer U (2016) SPATA2 promotes CYLD activity and regulates TNF-induced NF-kappaB signaling and cell death. EMBO Rep 17: 1485–1497

Shin D, Mukherjee R, Liu Y, Gonzalez A, Bonn F, Liu Y, Rogov VV, Heinz M, Stolz A, Hummer G, Dotsch V, Luo ZQ, Bhogaraju S, Dikic I (2020) Regulation of Phosphoribosyl-Linked Serine Ubiquitination by Deubiquitinases DupA and DupB. Mol Cell 77: 164–179 e6

Siegel LM, Monty KJ (1966) Determination of molecular weights and frictional ratios of proteins in impure systems by use of gel filtration and density gradient centrifugation. Application to crude preparations of sulfite and hydroxylamine reductases. Biochimica et Biophysica Acta (BBA) - Biophysics including Photosynthesis 112: 346–362

Smit JJ, Monteferrario D, Noordermeer SM, van Dijk WJ, van der Reijden BA, Sixma TK (2012) The E3 ligase HOIP specifies linear ubiquitin chain assembly through its RING-IBR-RING domain and the unique LDD extension. The EMBO Journal 31: 3833–3844

Smit JJ, van Dijk WJ, El Atmioui D, Merkx R, Ovaa H, Sixma TK (2013) Target specificity of the E3 ligase LUBAC for ubiquitin and NEMO relies on different minimal requirements. J Biol Chem 288: 31728–37

Sonn-Segev A, Belacic K, Bodrug T, Young G, VanderLinden RT, Schulman BA, Schimpf J, Friedrich T, Dip PV, Schwartz TU, Bauer B, Peters J-M, Struwe WB, Benesch JLP, Brown NG, Haselbach D, Kukura P (2019) Quantifying the heterogeneity of macromolecular machines by mass photometry. bioRxiv: 864553

Stieglitz B, Morris-Davies AC, Koliopoulos MG, Christodoulou E, Rittinger K (2012) LUBAC synthesizes linear ubiquitin chains via a thioester intermediate. EMBO Rep 13: 840–6

Stieglitz B, Rana RR, Koliopoulos MG, Morris-Davies AC, Schaeffer V, Christodoulou E, Howell S, Brown NR, Dikic I, Rittinger K (2013) Structural basis for ligase-specific conjugation of linear ubiquitin chains by HOIP. Nature 503: 422–426

Swatek KN, Usher JL, Kueck AF, Gladkova C, Mevissen TET, Pruneda JN, Skern T, Komander D (2019) Insights into ubiquitin chain architecture using Ub-clipping. Nature 572: 533–537

Taus T, Kocher T, Pichler P, Paschke C, Schmidt A, Henrich C, Mechtler K (2011) Universal and confident phosphorylation site localization using phosphoRS. J Proteome Res 10: 5354–62

Tokunaga F, Nakagawa T, Nakahara M, Saeki Y, Taniguchi M, Sakata S, Tanaka K, Nakano H, Iwai K (2011) SHARPIN is a component of the NF-kappaB-activating linear ubiquitin chain assembly complex. Nature 471: 633–6

Tokunaga F, Sakata S-i, Saeki Y, Satomi Y, Kirisako T, Kamei K, Nakagawa T, Kato M, Murata S, Yamaoka S, Yamamoto M, Akira S, Takao T, Tanaka K, Iwai K (2009) Involvement of linear polyubiquitylation of NEMO in NF-κB activation. Nature Cell Biology 11: 123–132

van Well EM, Bader V, Patra M, Sanchez-Vicente A, Meschede J, Furthmann N, Schnack C, Blusch A, Longworth J, Petrasch-Parwez E, Mori K, Arzberger T, Trumbach D, Angersbach L, Showkat C, Sehr DA, Berlemann LA, Goldmann P, Clement AM, Behl C et al. (2019) A protein quality control pathway regulated by linear ubiquitination. EMBO J 38

van Wijk SJL, Fricke F, Herhaus L, Gupta J, Hotte K, Pampaloni F, Grumati P, Kaulich M, Sou YS, Komatsu M, Greten FR, Fulda S, Heilemann M, Dikic I (2017) Linear ubiquitination of cytosolic Salmonella Typhimurium activates NF-kappaB and restricts bacterial proliferation. Nat Microbiol 2: 17066

Wagner S, Carpentier I, Rogov V, Kreike M, Ikeda F, Lohr F, Wu CJ, Ashwell JD, Dotsch V, Dikic I, Beyaert R (2008) Ubiquitin binding mediates the NF-kappaB inhibitory potential of ABIN proteins. Oncogene 27: 3739–45

Wagner SA, Satpathy S, Beli P, Choudhary C (2016) SPATA2 links CYLD to the TNF-alpha receptor signaling complex and modulates the receptor signaling outcomes. EMBO J 35: 1868–84

Yagi H, Ishimoto K, Hiromoto T, Fujita H, Mizushima T, Uekusa Y, Yagi-Utsumi M, Kurimoto E, Noda M, Uchiyama S, Tokunaga F, Iwai K, Kato K (2012) A non-canonical UBA-UBL interaction forms the linear-ubiquitin-chain assembly complex. EMBO Rep 13: 462–8

Ye Y, Akutsu M, Reyes-Turcu F, Enchev RI, Wilkinson KD, Komander D (2011) Polyubiquitin binding and cross-reactivity in the USP domain deubiquitinase USP21. EMBO Rep 12: 350–7

Young G, Hundt N, Cole D, Fineberg A, Andrecka J, Tyler A, Olerinyova A, Ansari A, Marklund EG, Collier MP, Chandler SA, Tkachenko O, Allen J, Crispin M, Billington N, Takagi Y, Sellers JR, Eichmann C, Selenko P, Frey L et al. (2018) Quantitative mass imaging of single biological macromolecules. Science 360: 423–427

Zivanov J, Nakane T, Forsberg BO, Kimanius D, Hagen WJ, Lindahl E, Scheres SH (2018) New tools for automated high-resolution cryo-EM structure determination in RELION-3. Elife 7

