## Supplementary material for "The linear ubiquitin chain assembly complex LUBAC generates heterotypic ubiquitin chains": EV Table 4

| gRNA1 forward | 5’-CACCGGTGTGGCAGACTGTACAG-3’ |
| --- | --- |
| gRNA1 reverse | 5’-AAACCTGTACAGTCTGCCACACC-3’ |
| gRNA63 forward | 5’-CACCGTGTGGTGCAGAAGAAGGA-3’ |
| gRNA63 reverse | 5’-AAACACACCACGTCTTCTTCCTC-3’ |
| HOIP mutagenesis C885A forward | 5’-AGGCGCCATGCACTTTCACTGTACCCAGTGCCGCCACCAG  -3’ |
| HOIP mutagenesis C885A reverse | 5’-GTGAAAGTGCATGGCGCCTCCTCGGGCCAGGGCGTACG-3’ |
| HOIL-1L mutagenesis C460A forward | 5’-AAGAAGGACGGCGCAGACTGGATCCGCTGCACCGTCTGCC-3’ |
| HOIL-1L mutagenesis C460A reverse | 5’-AGTCTGCGCCGTCCTTCTTCTGTACCACGATCTGGCACTGGGGG-3’ |
| HOIL-1L mutagenesis T203A/R210A forward | 5’-CTTCATCAACAAGCCCACGGCGCCTGGCTGTGAGATGTGCTGC-3’ |
| HOIL-1L mutagenesis T203A/R210A reverse | 5’-CCGTGGGCTTGTTGATGAAGGCGCACCCGGGGCACTGCCAGCC-3’ |
| ITA fragment generation backbone forward | 5’-GGTGGGTAACCTAGGCTG-3’ |
| ITA fragment generation backbone reverse | 5’-CGAAGATCTGCATGGTATATCTCCTTCTTAAAG-3’ |
| ITA fragment generation linker forward | 5’-GAGGTGGGTAAGAATTCGAG-3’ |
| ITA fragment generation linker reverse | 5’-CACGAAGATCTGCATATGTATATCTCCTTCTTATACTTAAC-3’ |
| ITA fragment generation ubiquitin 1 forward | 5’-GAAGGAGATATACCATGCAGATCTTCGTGAAGACCCTG-3’ |
| ITA fragment generation ubiquitin 1 reverse | 5’-CGCCGAGCTCGAATTCTTACCCACCTCTCAGGCGAAG-3’ |
| ITA fragment generation ubiquitin 2 forward | 5’-GAAGGAGATATACATATGCAGATCTTCGTGAAGACCCTG-3’ |
| ITA fragment generation ubiquitin 2 reverse | 5’-GCCTAGGTTACCCACCTCTCAGGCGAAG-3’ |
| Ubiquitin T12V mutagenesis forward | 5’-GACTGGCAAGGTCATCACCCTGGAGG-3’ |
| Ubiquitin T12V mutagenesis reverse | 5’-GGTGATGACCTTGCCAGTCAGGGTC-3’ |
| Ubiquitin T55V mutagenesis forward | 5’-GATGGCCGCGTCCTCTCTGATTAC-3’ |
| Ubiquitin T55V mutagenesis reverse | 5’-GAGAGGACGCGGCCATCTTCCAG-3’ |

**EV Table 1 Primers**
