## Supplemental Figures for "The linear ubiquitin chain assembly complex LUBAC generates heterotypic ubiquitin chains"

**Figure EV1 – Gel filtration analysis of LUBAC showing presence of multiple populations with different oligomeric states.**

**A** Gel filtration profile of purified LUBAC separated over S200 column.

**B** Tandem gel filtration separation of fraction 3 re-run over S200 column.

**C** Molecular weight standards separated over S200 column.

**Figure EV2 – Modelling of the LUBAC complex by negative staining electron microscopy.**

**A** LUBAC 2D class averages obtained from negatively stained particles.

**B** Initial 3D model of LUBAC made from particles picked in negatively stained electron micrographs.

**Figure EV3 – Projections made from 3D refined model of LUBAC.**

**Figure EV4 – Independent mass photometry measurements of LUBAC.**

**Figure EV5 – Purification of LUBAC complexes containing catalytically inert HOIP and HOIL-1L proteins.**

**A** SDS-PAGE analysis of different purified LUBAC complexes.

**B** Immunoblot analysis of different purified LUBAC complexes.

**Figure EV6 – Generation of *Hoil-1l*$^{C458A/C458A}$ mice.**

**A** Sequences of genomic DNA around C458 codon, gRNAs, and donor oligonucleotide used to target HOIL-1L C458A mutation.

**B** Sanger sequencing confirming correct mutations at target sites.

**C** Genotyping results of *Hoil-1l*$^{+/+}$, *Hoil-1l*$^{+/C458A}$, and *Hoil-1l*$^{C458A/C458A}$ mice. Hpy188III digest of a PCR fragment confirming correct targeting where a silent mutation is inserted.

**D** Immunoblot analysis of LUBAC component expression in MEFs derived from *Hoil-1l*$^{+/+}$ and *Hoil-1l*$^{C458A/C458A}$ mice.

**Figure EV7 – Anti-linear ubiquitin antibody validation.**

**A** Detection of mono-ubiquitin and di-ubiquitin of different linkages by anti linear ubiquitin antibody.

**B** Detection of longer Lys48 and Lys63-linked ubiquitin chains by anti-linear ubiquitin antibody.

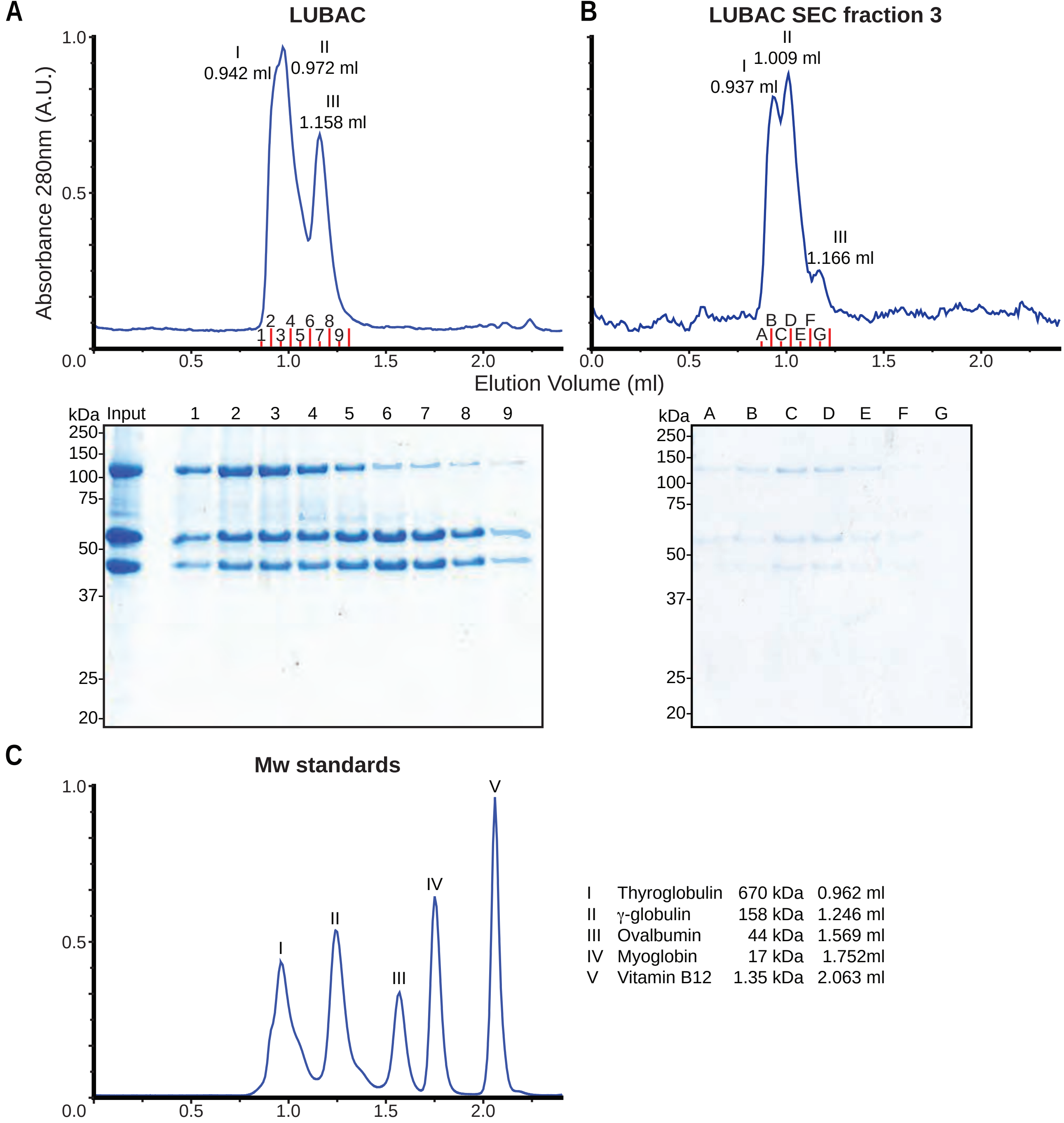
A
LUBAC
Absorbance 280nm (A.U.)
I 0.942 ml
II 0.972 ml
III 1.158 ml
B
LUBAC SEC fraction 3
I 0.937 ml
II 1.009 ml
III 1.166 ml
Elution Volume (ml)
kDa Input 1 2 3 4 5 6 7 8 9
250 150 100 75 50 37 25 20
kDa A B C D E F G
C
Mw standards
I Thyroglobulin 670 kDa 0.962 ml
II γ-globulin 158 kDa 1.246 ml
III Ovalbumin 44 kDa 1.569 ml
IV Myoglobin 17 kDa 1.752ml
V Vitamin B12 1.35 kDa 2.063 ml

**Figure EV1**

A

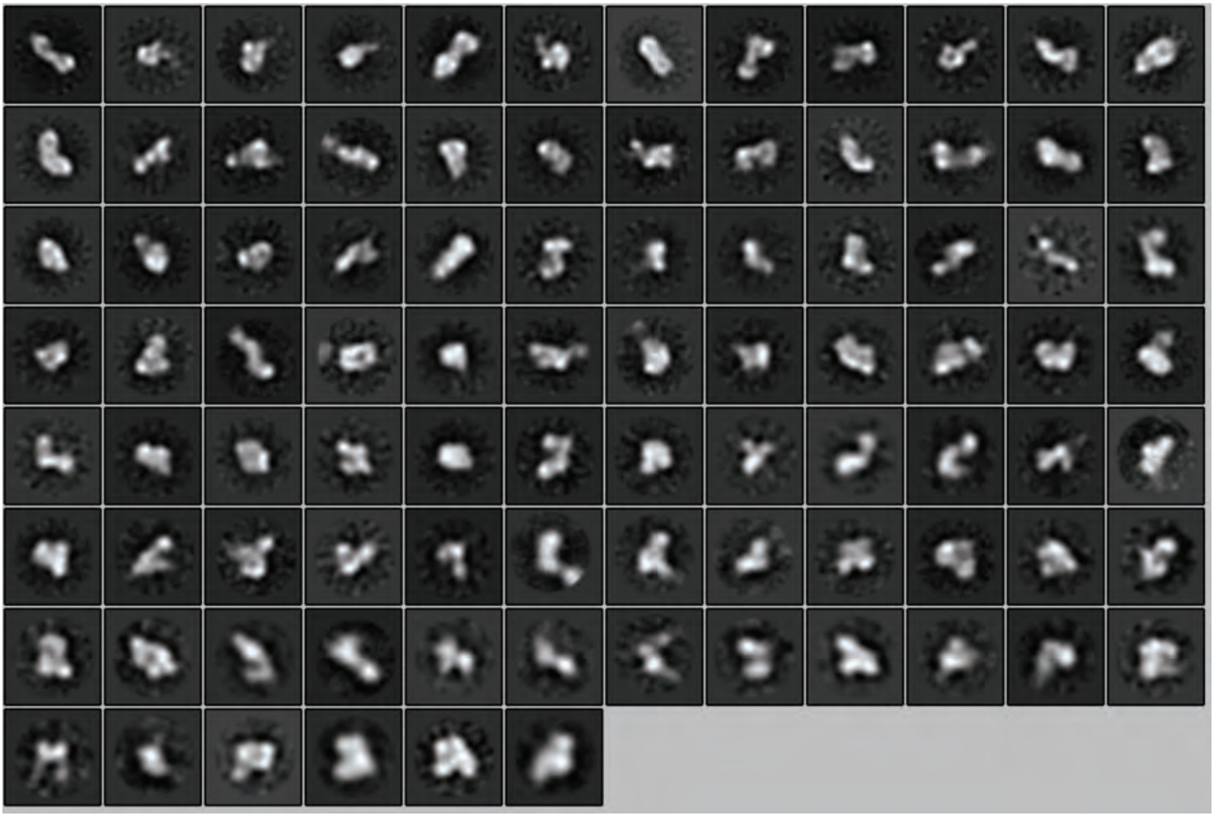

B

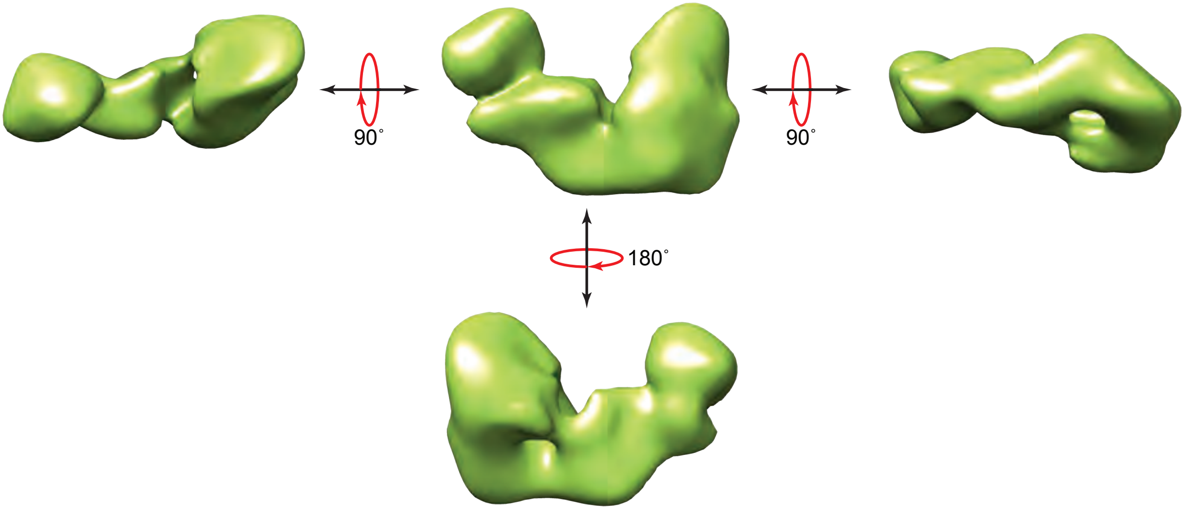


Figure EV2

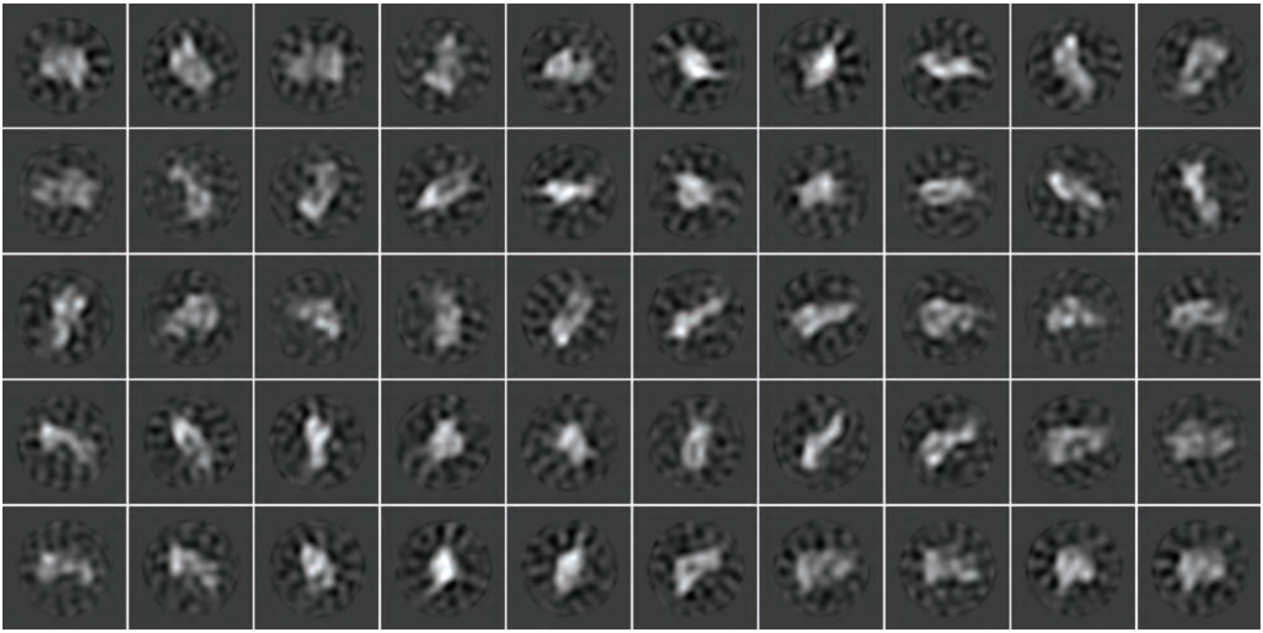

Figure EV3

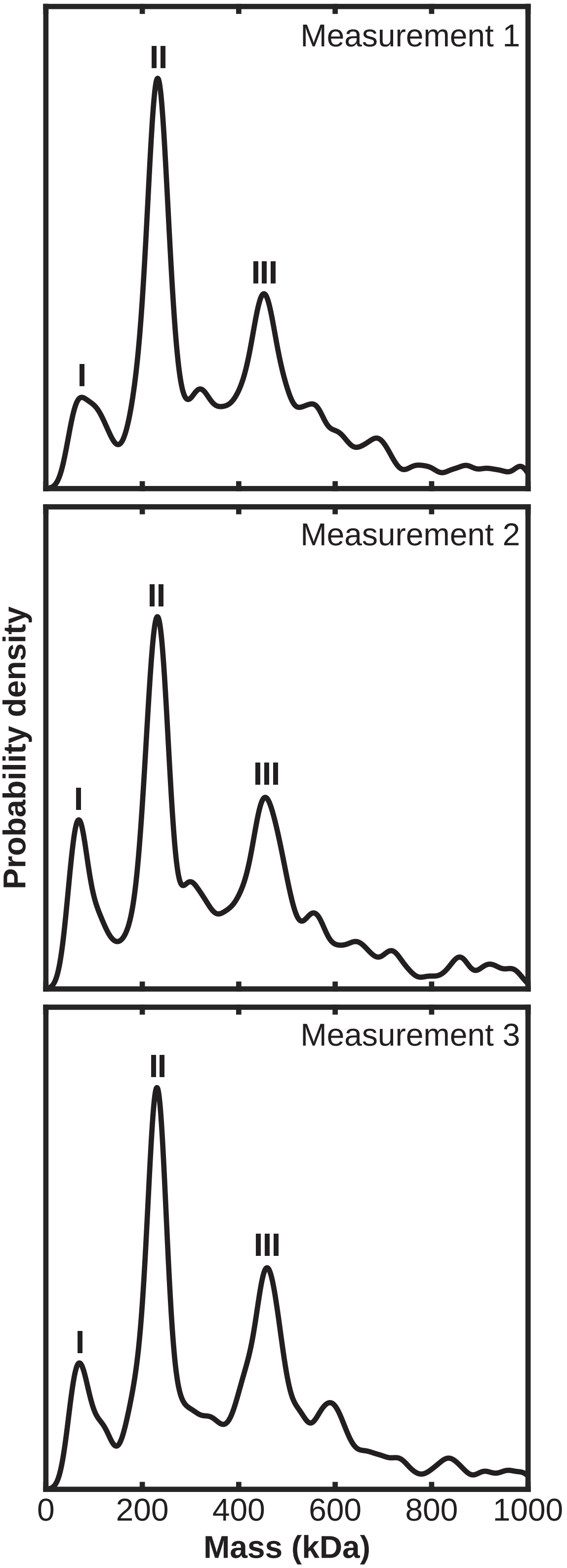

Measurement 1
I
II
III
Measurement 2
I
II
III
Measurement 3
I
II
III
Probability density
0
200
400
600
800
1000
Mass (kDa)


**Figure EV4**

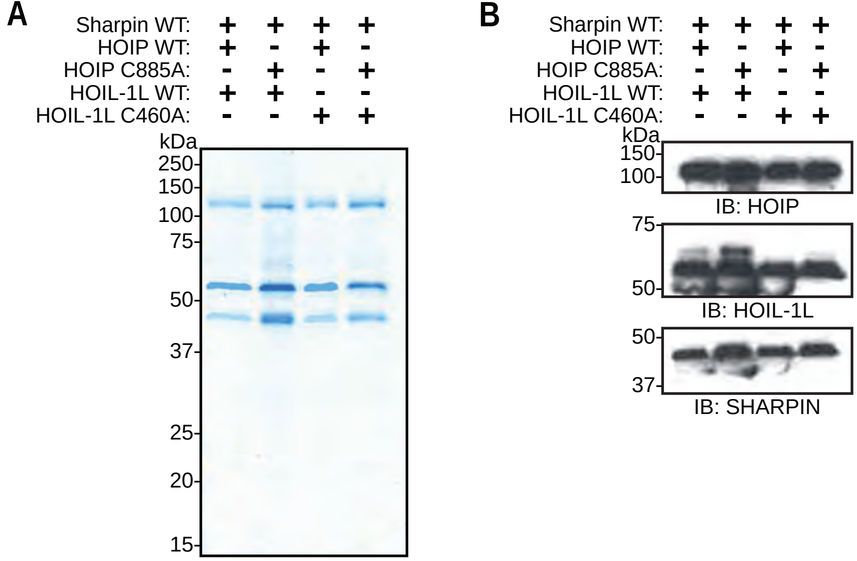
A
Sharpin WT: + + + +
HOIP WT: + - + -
HOIP C885A: - + - +
HOIL-1L WT: + + - -
HOIL-1L C460A: - - + +
kDa
250
150
100
75
50
37
25
20
15
B
Sharpin WT: + + + +
HOIP WT: + - + -
HOIP C885A: - + - +
HOIL-1L WT: + + - -
HOIL-1L C460A: - - + +
kDa
150
100
IB: HOIP
75
50
IB: HOIL-1L
50
37
IB: SHARPIN

Figure EV5

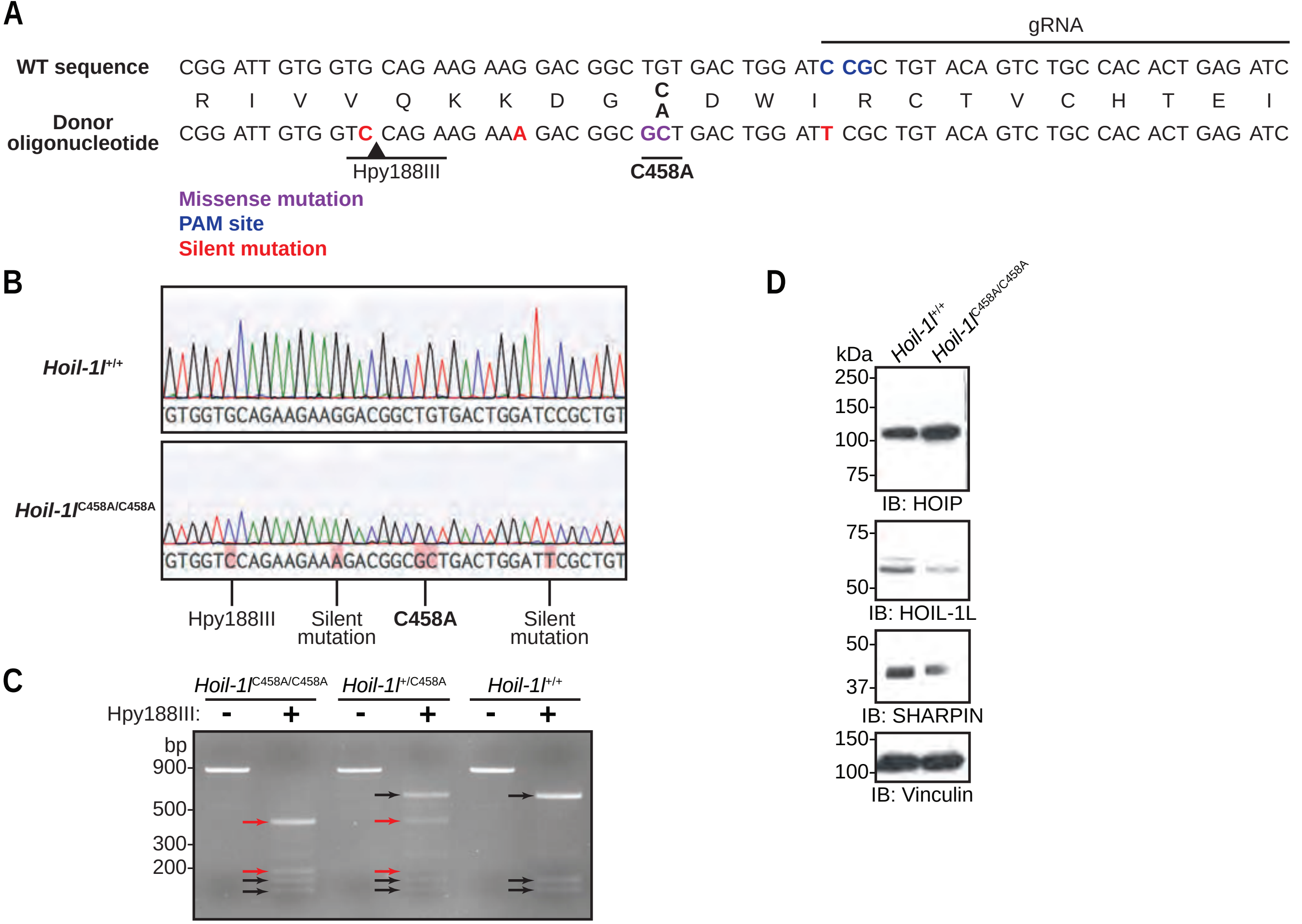
A
gRNA
WT sequence
CGG ATT GTG GTG CAG AAG AAG GAC GGC TGT GAC TGG ATC CGC TGT ACA GTC TGC CAC ACT GAG ATC
R I V V Q K K D G C A D W I R C T V C H T E I
Donor oligonucleotide
CGG ATT GTG GTC CAG AAG AAA GAC GGC GCT GAC TGG ATT CGC TGT ACA GTC TGC CAC ACT GAG ATC
Hpy188III
C458A
Missense mutation
PAM site
Silent mutation
B
Hoil-1l+/+
GTGGTGCAGAAGAAGGACGGCTGTGACTGGATCCGCTGT
Hoil-1lC458A/C458A
GTGGTCCAGAAGAAAGACGGCGCTGACTGGATTCGCTGT
Hpy188III
Silent mutation
C458A
Silent mutation
C
Hoil-1lC458A/C458A
Hoil-1l+/C458A
Hoil-1l+/+
Hpy188III: - + - + - +
bp
900-
500-
300-
200-
D
Hoil-1l+/+
Hoil-1lC458A/C458A
kDa
250-
150-
100-
75-
IB: HOIP
75-
50-
IB: HOIL-1L
50-
37-
IB: SHARPIN
150-
100-
IB: Vinculin

Figure EV6

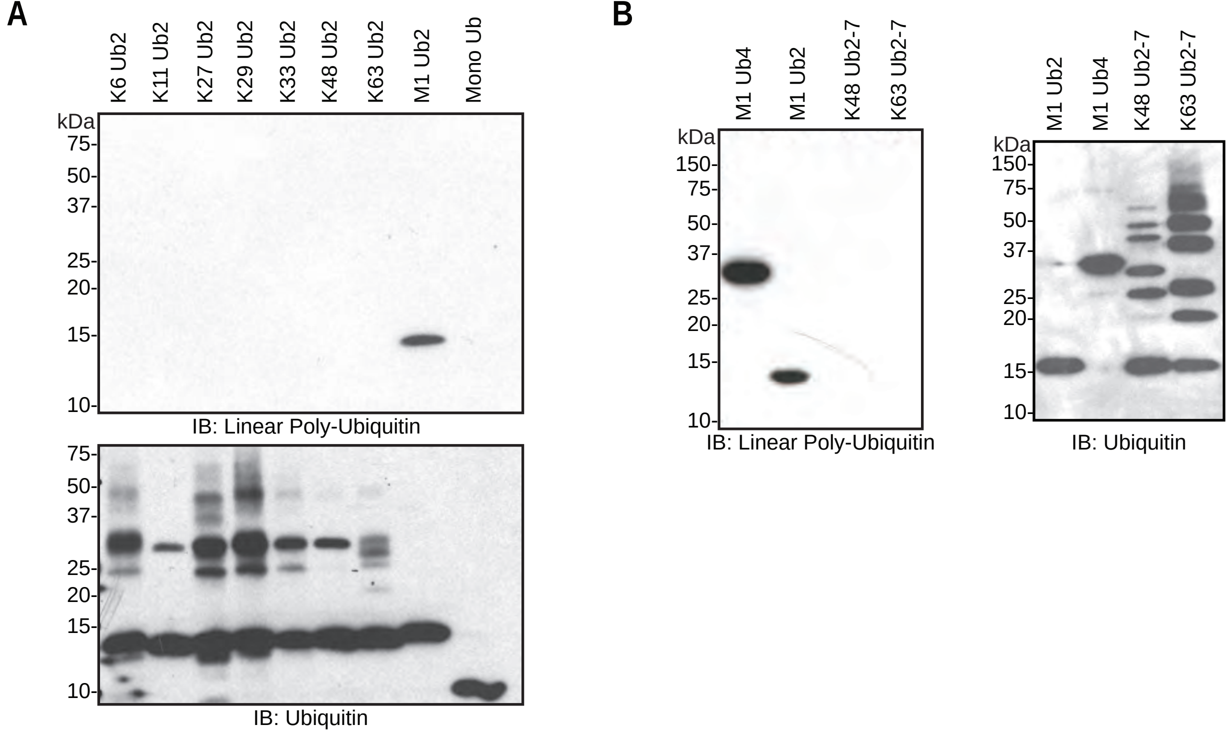
A
K6 Ub2
K11 Ub2
K27 Ub2
K29 Ub2
K33 Ub2
K48 Ub2
K63 Ub2
M1 Ub2
Mono Ub
kDa
75-
50-
37-
25-
20-
15-
10-
IB: Linear Poly-Ubiquitin
75-
50-
37-
25-
20-
15-
10-
IB: Ubiquitin
B
M1 Ub4
M1 Ub2
K48 Ub2-7
K63 Ub2-7
kDa
150-
75-
50-
37-
25-
20-
15-
10-
IB: Linear Poly-Ubiquitin
M1 Ub2
M1 Ub4
K48 Ub2-7
K63 Ub2-7
kDa
150-
75-
50-
37-
25-
20-
15-
10-
IB: Ubiquitin

**Figure EV7**
